# Local Temperature and Humidity are Associated with Proportion of Antimicrobial-Resistant *Escherichia coli* isolates in Farm Environments: Considerations for On-Farm Surveillance

**DOI:** 10.64898/2026.05.27.728203

**Authors:** Lucy Vass, Elliot Stanton, Hannah Schubert, Katy Morley, Emma F. Puddy, Fernando Sánchez-Vizcaíno, Virginia C. Gould, Oliver Mounsey, Matthew B. Avison, Kristen K. Reyher, Andrew W. Dowsey

## Abstract

Evidence suggests that increased local temperatures are associated with higher prevalence of antimicrobial resistance (AMR) in environmental bacteria. This study investigates the association between local climate and the proportion of antimicrobial-resistant *Escherichia coli* isolated from 2,766 farm environment samples from 53 English dairy farms. To do this, a non-linear Bayesian model that specifically accounts for decreased test sensitivity at low *E. coli* abundance was developed and used to estimate the proportion of isolates resistant to four antimicrobials (amoxicillin, cephalexin, streptomycin and tetracycline) from colony count data. Mean 7-day temperature and relative humidity at the farm location was modelled using a generalised additive model formulation. A higher proportion of *E. coli* isolates were resistant to cephalexin and streptomycin in samples collected from adult cow collecting yards, than heifer housing sheds. In contrast, a greater proportion of *E. coli* isolates from heifer housing sheds were resistant to amoxicillin and tetracycline. Evidence that local temperature is associated with an increase in the proportion of *E. coli* isolates resistant to streptomycin (20°C increase associated with a 5.0-fold increase; 95% CI: 1.03-33.0) and tetracycline (2.6-fold increase; 90% CI: 1.1-5.2) was observed. Additionally, relative humidity was associated with an increase in the proportion of isolates resistant to amoxicillin streptomycin and tetracycline. The influence of weather on the proportion of antimicrobial-resistant *E. coli* varied between samples collected from adult animals in collecting yards and heifers in housing sheds. These findings highlight the importance of considering weather conditions, sample characterises and seasonality when designing on-farm AMR surveillance systems.

**Importance:** Understanding how environmental conditions are associated with variability in AMR prevalence is critical for developing robust livestock AMR surveillance and anticipating the potential effects of climate change. The non-linear Bayesian modelling approach developed here adjusts for *E. coli* abundance associated variability in test sensitivity, enabling the influence of risk factors associated with the proportion of antimicrobial-resistant *E. coli* within samples to be more accurately estimated. Applying this approach to 2,766 faecal samples from 53 dairy farms in Southwest England indicated that the proportion of antimicrobial-resistant *E. coli* generally increased under warmer and wetter conditions. These findings suggest that environmental conditions can influence the prevalence of AMR *E. coli* in dairy farm environments and demonstrate the importance of accounting for weather related variability in livestock AMR surveillance. Adjusting for these associations in livestock AMR surveillance could improve the accuracy of modelling AMR trends and strengthen the assessment of climate-associated AMR risks.

## Introduction

Antimicrobial resistance (AMR) and climate change are considered key threats to global public health by many, including the World Health Organisation (1). In addition to the threat they each pose separately, climate change could amplify and accelerate AMR through increased bacterial infections, food insecurity and increased overlap of the environments between humans, livestock and wildlife (2, 3). The climate and weather also play a role in the prevalence and dissemination of AMR genes (ARGs). It is known that warmer temperatures relevant to UK weather conditions are associated with increased bacterial growth rate and survival, increased rate of horizontal gene transfer (HGT; 4, 5) and improved fitness of antimicrobial-resistant *E. coli* at high temperatures due to the co-selection of ARGs with stress response genes (6, 7).

Understanding the relationship between climate and AMR is essential for estimating the burden of AMR and potential geographical variation in future years, and hence key to selection and prioritisation of possible interventions. Current predictions - do not consider this effect and could therefore be underestimates (8, 9).

There is a range of evidence emerging which indicates that, at the population level, humans living in warmer climates have a higher prevalence of important AMR bacterial infections. One such ecological study, by MacFadden *et al.*, (2018) assessed the rate of AMR in three important human pathogens (*Escherichia coli, Klebsiella pneumoniae* and *Staphylococcus aureus*) and found a positive association with 30-year mean minimum temperature local temperature. The effect size seen by these authors was greatest in *E. coli,* where a 10°C increase in temperature was associated with an average increase of 4.2% in the proportion of isolates with any AMR (*p* < 0.0001; 10). Similar ecological studies in Europe have mirrored these results with positive associations between country-level annual average minimum temperature and AMR in all classes of antimicrobials and pathogens tested (11, 12).

The environment is recognised serve as key compartment in One Health AMR, and provides an interface between humans, livestock and wildlife (13). Positive associations between temperature and AMR have also been demonstrated in the environmental microbiome. Temperature, for instance, is known to modulate species composition in the soil microbiome, and genomic analysis has revealed that the proportion and absolute quantity of ARGs within the soil microbiome increases with local temperature (14). This result has been mirrored in other environmental microbiomes, such as those of wetlands and rivers (15, 16).

The effect of temperature on AMR in food-producing animal populations has yet to be fully explored. Antimicrobial use in food-producing animals has come under scrutiny as evidence emerges suggesting that antimicrobial use and the development of AMR in these populations is related to AMR in human populations (17, 18). Aside from antimicrobial use, risk factors for the development of AMR within food-animal populations are poorly understood. A previous study by our group aiming to identify factors associated with the likelihood of detecting antimicrobial-resistant *E. coli* within the environment of UK dairy farms found a positive association with mean monthly temperature (19). In particular, the likelihood of detecting a mobile genetic element encoding resistance to 3^rd^ generation cephalosporins, CTX-M, was found to be increased at warmer mean monthly temperatures (OR: 1.57, 95% confidence interval: 1.20 - 2.06; 19).

The effect of temperature on AMR is also important to consider in the context of designing observational studies or surveillance systems, as changes in temperature and climatic conditions could lead to large seasonal changes in AMR prevalence (19). In addition to potentially impacting directly on AMR prevalence, temperature changes can impact the detection of AMR through the effect of temperature on bacterial survival, growth and abundance (20). If bacterial abundances within samples are not standardised, or if the microbiology techniques used do not account for differing bacterial abundances across timepoints, the likelihood of detecting AMR bacteria could vary greatly between seasons, impacting the results of observational studies or surveillance.

This study investigates local weather conditions and their interactions as potential risk factors for antimicrobial-resistant *E. coli* using data collected as part of Schubert *et al.* (2021). A novel, non-linear Bayesian model with a Poisson likelihood was developed, allowing appropriate adjustment for sample-level heterogeneity in *E. coli* abundance. This approach is well suited to livestock AMR surveillance contexts as it accounts for bacterial abundance-associated variability in test sensitivity which could obscure associations between potential risk factors such as local weather conditions and the proportion of antimicrobial-resistant *E. coli* within samples.

## Materials and methods

### Sample collection and microbiology

A convenience sample of 53 dairy cattle farms in the southwest region of England, UK, were recruited and followed for 2-years (detailed in 20). A subset of faecally contaminated environmental which were collected from either: the collecting yard (termed AY; an often-uncovered yard where adult milking cattle wait before entering the milking parlour, n = 1433) or from heifer sheds (termed HS; typically roof-covered housing for female weaned cattle before their first calving, n = 1333) between January 2017 and July 2018 were considered within this study. To collect samples, a researcher traversed the site wearing sterile plastic overboot socks (method adapted from 21). To prepare for antimicrobial sensitivity testing, the faecally contaminated material on the overshoes was homogenised in 10mL/g of phosphate-buffered saline (PBS). Samples were suspended in 50% sterile glycerol (v/v) (1:1) and an aliquot was stored at −80°C before culture. A 20-microliter aliquot of the sample was diluted 10-fold and cultured on agar (Tryptone bile x-glucuronide (TBX); Scientific Laboratory Supplies). A further 4 undiluted 20-microlitre aliquots were taken and cultured on TBX agar containing one of 5 antimicrobials at concentrations relevant to resistance in humans according to EUCAST clinical breakpoints (amoxicillin at 8mg/L, cephalexin at 16mg/L, streptomycin at 16mg/L, tetracycline at 16mg/L and ciprofloxacin at 0.5 mg/L; 22). After incubation at 37°C (overnight), the number of *E. coli* colony-forming units (CFU) was then manually counted on each plate to give an estimate of *E. coli* abundance in the sample and the proportion of *E. coli* resistant to each antimicrobial tested (Figure 1).

**Figure 1:**
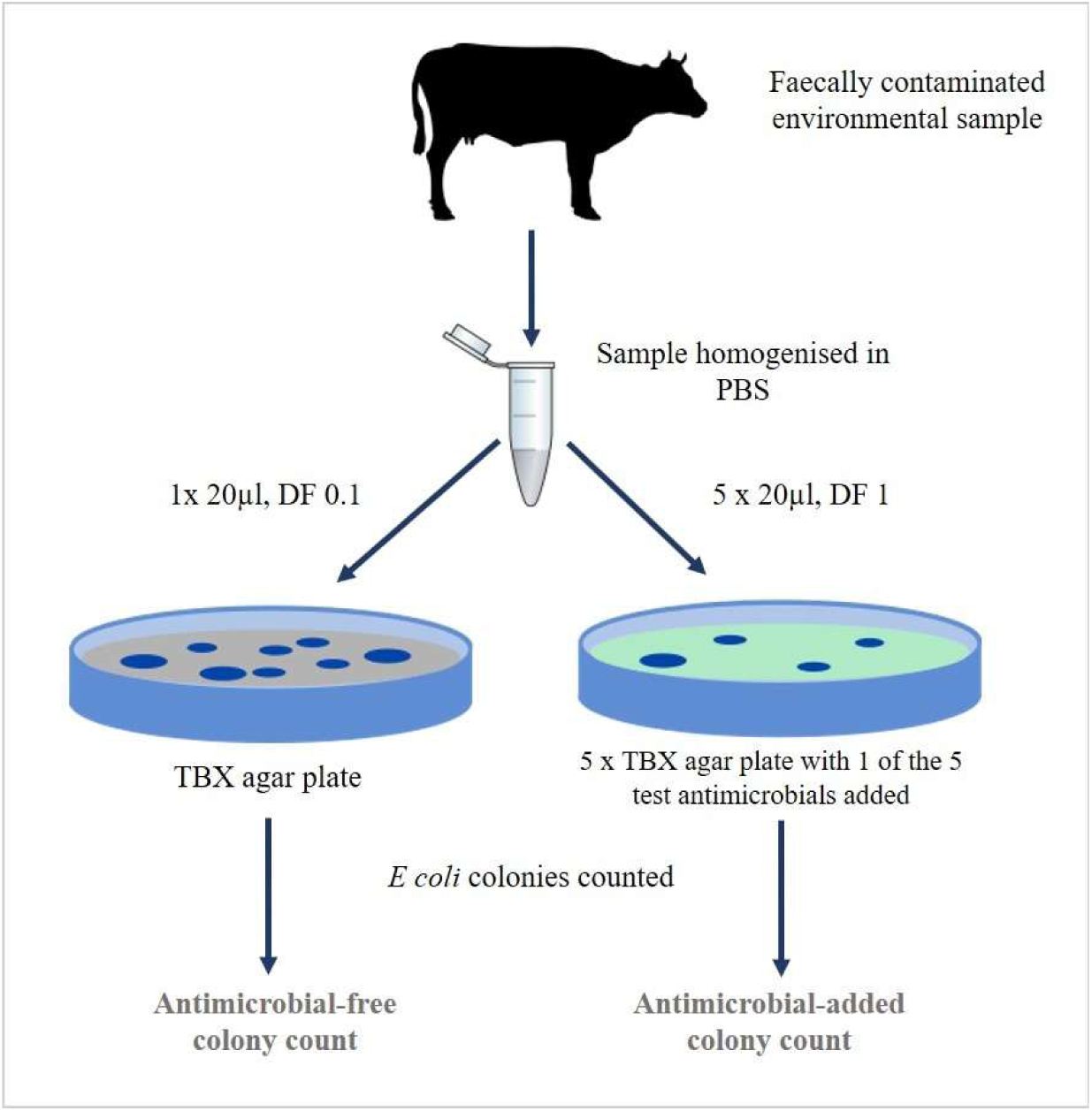
Sample plating and antimicrobial sensitivity testing. *Diagram showing the microbiology methodology for antimicrobial sensitivity testing*. *DF: dilution factor. PBS: phosphate-buffered saline. TBX: tryptone bile X-glucuronide agar*.

Where plates had CFUs of >300, the CFU was estimated by counting a sub-sectioning of the plate. Samples where no *E. coli* colonies were detected on the AM-free plate were excluded from the analysis as missing data points (n = 173). These samples occurred across a range of climatic conditions (Figure S1).

### Meteorological data

Hourly air temperature (°C) and relative humidity (%) measurements were obtained from the Met Office Integrated Data Archive System (23). Each farm was matched to the nearest weather recording station by calculating the distance between each farm and the weather station using the haversine formula. The maximum distance between a study farm and the nearest station was 30km. Sample collection was assumed to be at noon and the preceding 7 days of hourly observations were used to calculate mean air temperature (Celsius) and mean relative humidity (%).

### Non-linear Bayesian model with Poisson likelihood for the proportion of antimicrobial-resistant isolates

The sample processing and method of detecting antimicrobial-resistant *E. coli* necessitated the use of a novel statistical model to estimate the proportion of resistant isolates within each sample. *E. coli* abundance within the samples varied significantly between sample location and over different temperatures (Figure S2). Low *E. coli* abundance was theorised to produce a higher limit of detection for AMR isolates as well as to decrease the accuracy and precision of an estimate of the proportion of antimicrobial-resistant *E. coli* within each sample (20). Statistical modelling was therefore required to capture the differing uncertainty produced by the varying *E. coli* abundance between samples. This ensured that we did not bias estimates of effects sizes of predictors potentially associated with both bacterial growth and proportion of AMR isolates, such as temperature and sample location.

A non-linear Bayesian model was developed based on a generalised linear model with Poisson likelihood. Two non-linear parameters were included in the model with sub-models set up to estimate these parameters. Sub-model 1 estimated a colony count for individual antimicrobial-free (AM-free) plates. Sub-model 2 estimated the proportion of resistant *E. coli* between each pair of AM-free and antimicrobial-added (AM-added) plates. The full model combined estimates of overall *E. coli* abundance and the proportion of antimicrobial-resistant *E. coli* to produce an estimated *E. coli* count for each agar plate. A diagrammatic representation of the non-linear Bayesian model with Poisson likelihood is shown in Figure S3. Prior specifications are detailed in Table S1 and justified in Figures S4 and S5.

To validate that the model produced realistic estimates of colony counts and proportions of antimicrobial-resistant *E. coli* while capturing variability as expected, model estimates from an intercept-only model were generated. Intercept-only models were first validated using a synthetic data set comprising a range of AM-free and AM-added colony counts (Figure S6). Following this, intercept-only models were applied to real data. Changes in uncertainty associated with *E. coli* abundance were first inspected, enabling comparison between models built using synthetic and real data (Figure S7). As had been done with the synthetic data (Figure S5), posterior predictive checks were conducted where the proportion of antimicrobial-resistant *E. coli* within a sample, estimated by dividing the AM-added count by the AM-free count, was compared to the model-estimated proportions of antimicrobial-resistant *E. coli* (Figure S8).

Performance was contrasted with that of an alternative model parameterisation using the zero-inflated Poisson distribution. Model comparison via posterior predictive checking (Figure S9) and expected log predictive density (ELPD; Table S2) was then performed. Evidence supporting the presence of excess zero counts was weak, with the mean estimate of the zero-inflation parameter being 0.027, suggesting that 2.7% of zero counts were in excess of those expected were the data Poisson distributed. Additionally, the ELPD of the simple Poisson model exceeded that of the zero-inflated Poisson model. This indicated that the original Poisson likelihood model was the more appropriate choice and so this model structure was carried forward.

All models were run in R using the Bayesian modelling package *brms* (24) using the *cmdstanr* backend (25). All models were run across 4 Markov Chain Monte Carlo (MCMC) chains each with 20,000 total sampling steps. The first 50% of draws were discarded as warm up draws, giving a total of 40,000 posterior draws. The convergence of the MCMC chains was checked using the diagnostic potential scale reduction factor (26). All models had sufficient burn-in and sampling iterations to reach a potential scale reduction factor of < 1.1 for parameters relating to the model predictors.

### Modelling the effect of weather on the proportion of antimicrobial-resistant *E. coli* (Sub-model 2)

A directed acyclic graph was constructed to visualise potential sample-level predictors for AMR in the farm environment as well as any correlations introduced by the study design (Figure 2).

**Figure 2:**
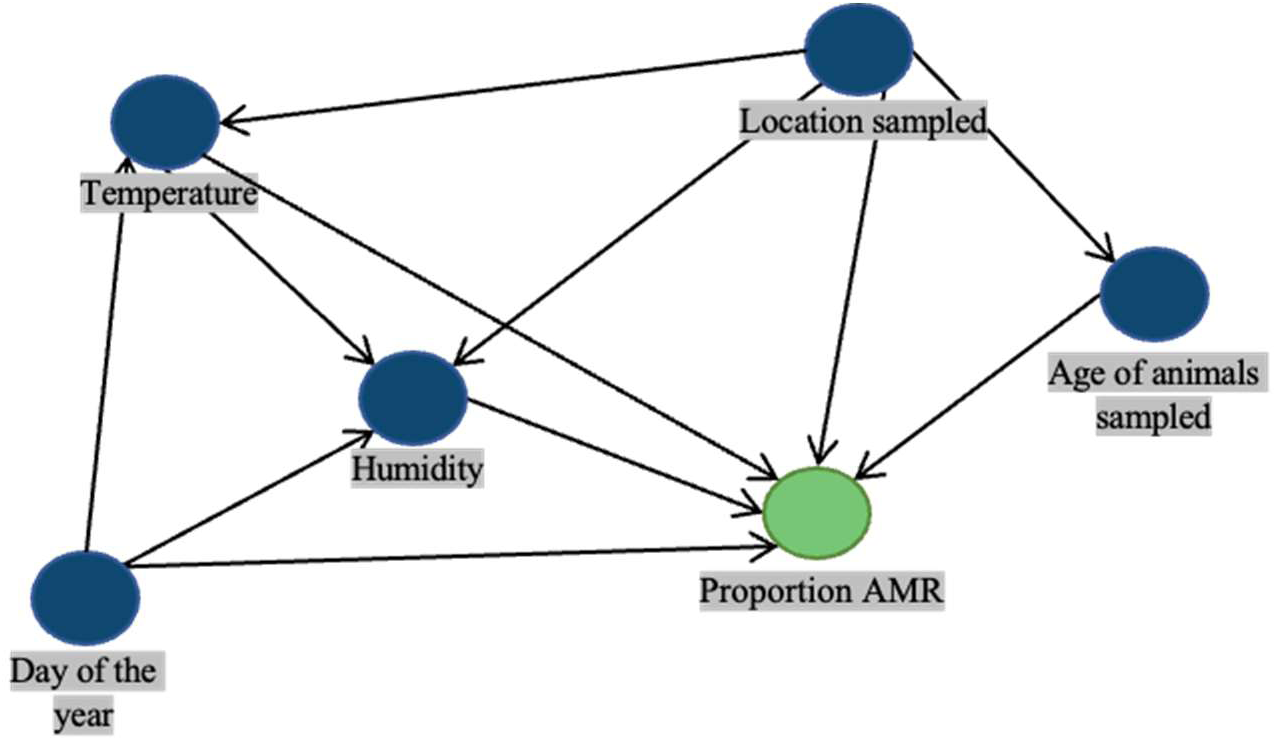
Directed acyclic graph showing potential causal relationships between exposures and the outcome, proportion of antimicrobial-resistant *E. coli* in the faecally contaminated environmental sample.

Aside from weather-related factors, the sampling location and age of animals populating the location were theorised to be associated with the proportion of antimicrobial-resistant *E. coli* isolates. As previous analysis of the complete dataset (including samples from pre-weaned calves) (19) found that samples from younger animals were associated with a higher likelihood of AMR, and other studies have evidenced similar associations in food-producing animals (27, 28), the effect of the age of cattle populating the sample site was considered. Sampling location was highly correlated with the age of cattle, as indoor housing was populated most often by heifers (weaned female cattle <2 years) and the collecting yard by adult milking cattle (>2 years). In addition, it was expected that outdoor local air temperature and humidity would be different to those at the sampling locations (unmeasured) due to cattle’s body heat, the insulating effects of buildings and reduced airflow. We therefore theorised that sampling location could modulate the effect of outdoor temperature and humidity on AMR. To account for these effects within the model, samples were split into two groups: AY (adult cattle in the collecting yard) and HS (heifers in sheds/pens) which were assessed in 2 separate models. This parameterisation was directly analogous to adding an interaction effect between each of the climatic predictors and age/location group. This resulted in 8 different outcome variables (AY and HS for each of the 4 antimicrobials).

We also considered other potential seasonal effects which were not explained by temperature or humidity, such as the differing seasonal management activities of the farm (‘Day of the year’ in Figure 2). These could include changes such as housing cattle indoors in the winter, lactation and calving periods as well as changes in disease prevalence and associated antimicrobial use. This ‘Day of the year’ effect was included in the multivariable models as a penalized thin plate regression spline using *brms’* support for the R package *mgcv* (29, 30, 31).

To account for repeated sampling from each location on the 53 farms, a nested group effect was added to the model. Each unique sampling location was nested within the corresponding farm.

### Assessing potential collinearity between temperature and relative humidity

Potential collinearity between the predictors for air temperature and relative humidity was assessed using Pearson’s correlation coefficient (with coefficients >0.7 or <-0.7 indicating potential collinearity). The effect size estimates for temperature and humidity were compared between univariable and multivariable (containing temperature and humidity) linear regression models to check for large changes in the direction or uncertainty of the effect size, either of which could indicate collinearity or another model misspecification.

### Non-linear effects for temperature and humidity

The association between temperature/humidity and the proportion of AMR isolates was theorised to be non-linear and was therefore modelled using penalized thin plate regression spline terms. A tensor-product smooth of the thin-plate regression splines was used to model a joint surface over temperature and humidity, allowing the combined effect of these covariates to vary smoothly across their observed ranges (29, 30, 31, 32). To assess if modelling the relationship using a tensor product smooth of thin plate regression splines improved model performance, the performance of a model containing linear effects for temperature and humidity was compared to models with the tensor product smooth via estimated log pointwise predictive density comparison. ELPD was estimated via k-fold cross validation using 5 folds using the R package *loo* (33).

### Model-estimated proportion of resistant isolates across different weather condition change scenarios

To quantitatively demonstrate the estimated effects of temperature and humidity on the proportion of antimicrobial-resistant *E. coli*, the model-estimated proportion of antimicrobial-resistant *E. coli* was compared between different local weather conditions. To avoid extrapolation, the values for temperature and humidity were chosen to be indicative of the minimum (1°C, 60% humidity), maximum (21°C, 92% humidity) and mean (10°C, 82% humidity) of the weather conditions encountered across the study period (when measured using 7-day mean). The change in the proportion of resistant isolates was estimated by the models across 4 weather condition change scenarios:

1. ‘Increase temperature’: Increase from 1°C to 21°C, at 82% humidity
2. ‘Increase humidity’: Increase from 60% to 92% humidity, at 10°C
3. ‘Positive interaction’: From cold & dry (1°C, 60%) to hot & humid (21°C, 92%)
4. ‘Negative interaction’: From cold & humid (1°C, 92%) to hot & dry (21°C, 60%).

The change in the estimated proportion of resistant isolates was calculated directly from all post-warm-up draws and summarised using median, 90% credible interval and 95% credible interval.

## Results

As few samples tested positive for ciprofloxacin resistant colonies, the estimated proportion of resistant isolates in these samples was generally very low (median: 2.90×10^-7^). Due to this and the lack of variance between samples, inferred effects for temperature and humidity predictors in uni- and multivariable ciprofloxacin models had high uncertainty. As a result, ciprofloxacin was not modelled in this study beyond the initial uni- and multivariable models.

Intercept-only models were used to estimate the proportion of antimicrobial-resistant *E. coli* colonies for each other antimicrobial. The highest estimated proportion of resistant *E. coli* colonies in all samples (AY and HS together) was found to the antimicrobial tetracycline (median: 1.8×10^-2^, 95% interval: 6.2×10^-4^ – 3.7×10^-1^) and the lowest to cephalexin (median: 2.62×10^-4^, 95% interval: 2.2×10^-5^ – 1.4×10^-1^). The estimated proportion of resistant colonies varied between sample location groups (samples taken from the collecting yard populated with adult dairy cattle [AY group, n = 1433] and samples taken from indoor sheds populated with heifers [HS group, n = 1333]). On average, the estimated proportions of colonies resistant to cephalexin and streptomycin were higher in the AY group than the HS group (AY median 39 times higher for cephalexin and 2.8 times higher for streptomycin; Figure 3). The opposite was true for amoxicillin and tetracycline, where the HS group had a higher estimated proportion of resistant colonies (HS median 1.8 times higher for amoxicillin and 5.1 times higher for tetracycline; Figure 3).

**Figure 3:**
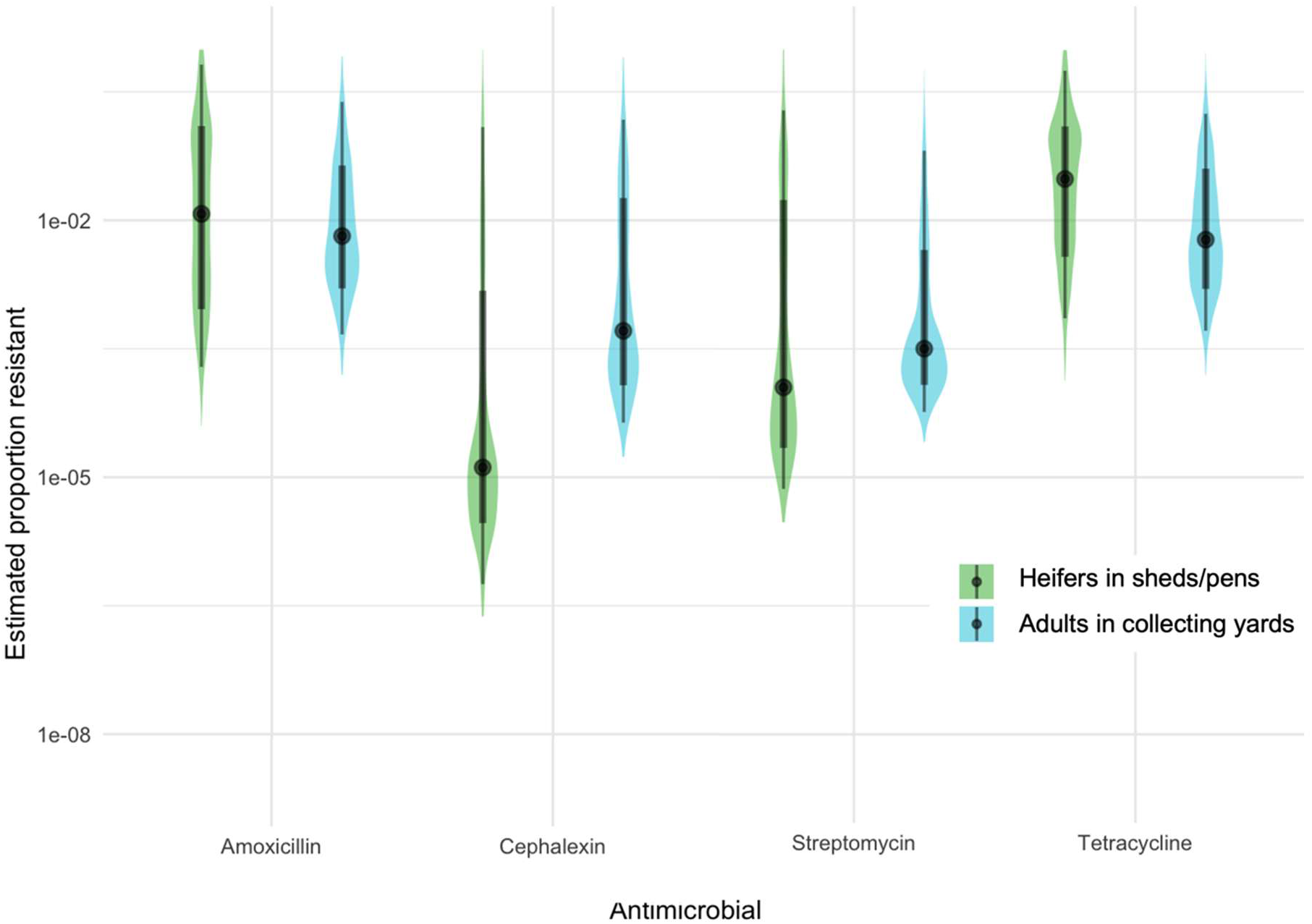
Distribution of estimated proportion of antimicrobial-resistant *E. coli* in all samples to each of the 4 tested antimicrobials. *Violin plots show the posterior distribution, black points, thick black bars and thin black bars show the median, 75% interval and 95% interval, respectively. The y-axis is log_10_ transformed. Estimated proportions were generated for each sample from an intercept only model*.

Across the 18-month sampling period considered, the mean 7-day air temperature was 10°C (min. 1°C, max. 21°C), and the mean 7-day relative humidity was 82% (min 60%, max 92%). When averaged over 7 days, temperature and humidity were weakly negatively correlated (Pearson’s correlation coefficient = −0.43). Comparing the effect size of temperature and humidity in univariable and multivariable models with linear effects for mean 7-day temperature, mean 7-day humidity and ‘day of study’ revealed no large changes in effect size, direction or uncertainty indicating there was no severe collinearity (Table S3).

### Multivariable model with splines for temperature and humidity

A multivariable model for the proportion of antimicrobial-resistant *E. coli* was constructed for each combination of sample group (HS and AY) and antimicrobial. Mean 7-day temperature and mean 7-day humidity were modelled as a tensor-product smooth of thin-plate regression splines and ‘day of study’ was modelled as a thin plate regression spline.

The results of these models were visualised to show the effect of temperature at a constant humidity, the effect of humidity at a constant temperature, and the interaction between temperature and humidity (Figure 4).

**Figure 4:**
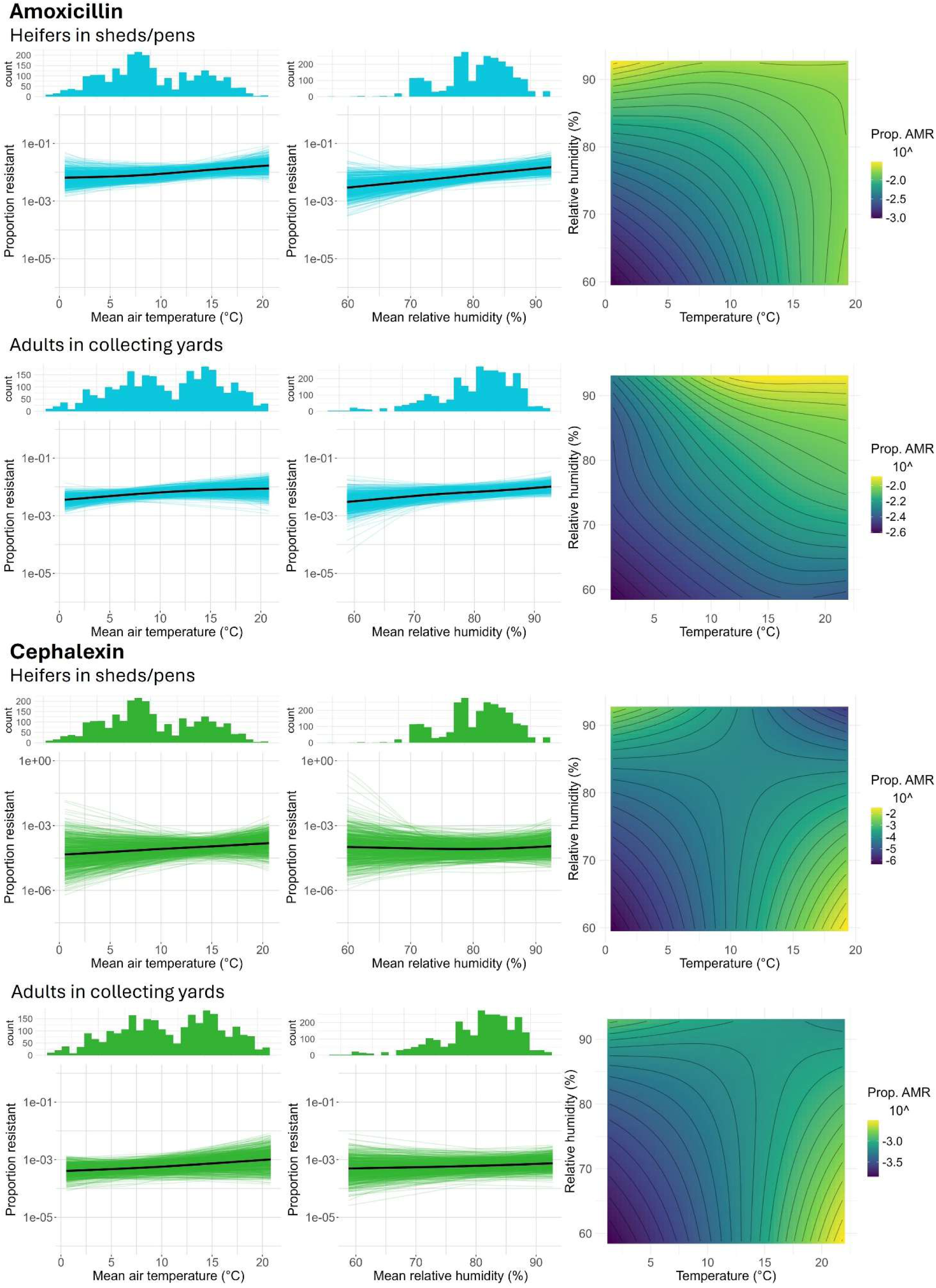
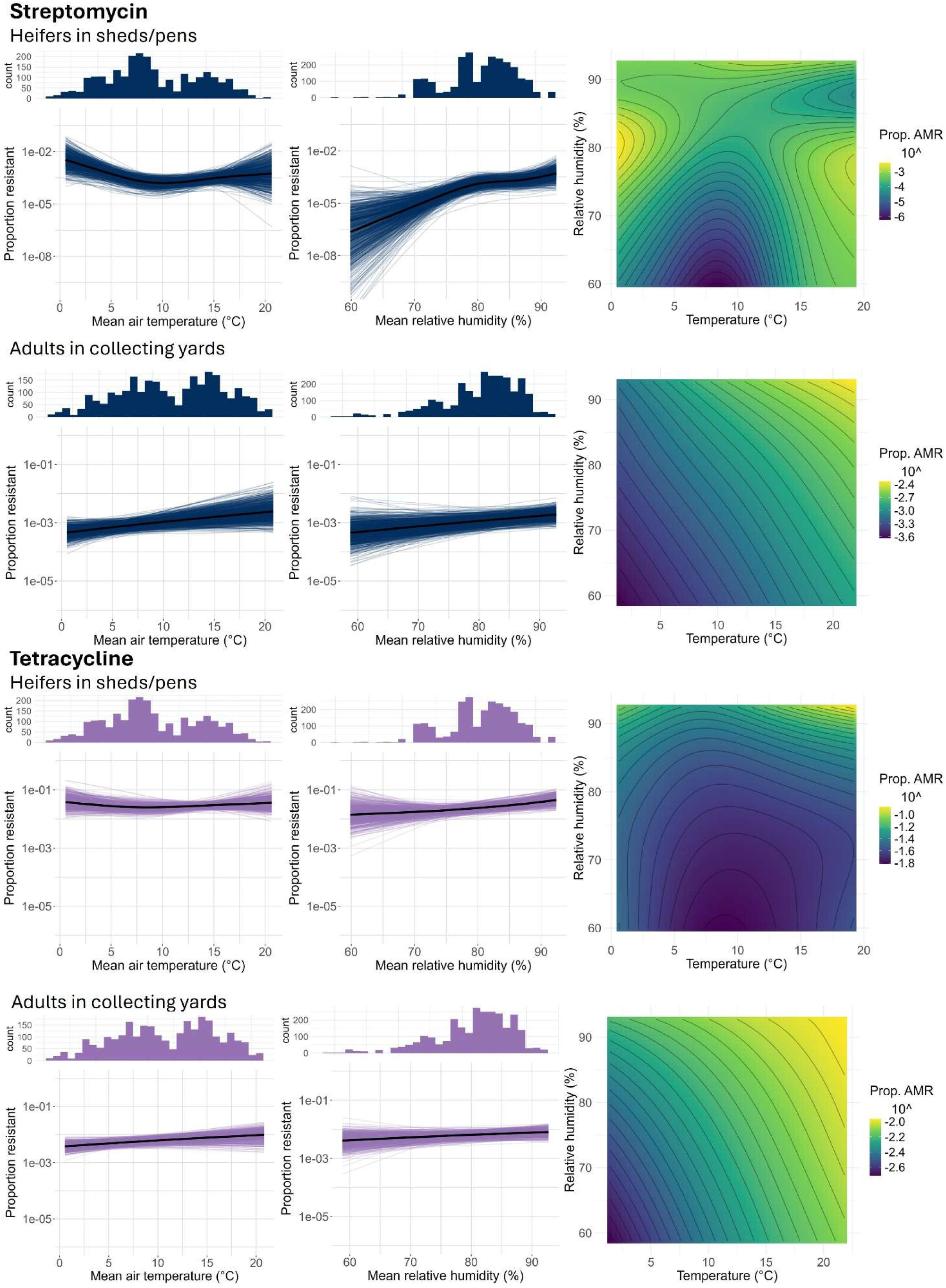
Estimated splines (left and centre columns) and tensor-product surfaces (right column) from non-linear models showing the effects of weather covariates on the estimated proportion of antimicrobial-resistant *E. coli* for each of the four tested antimicrobials across the two farm environments. *Splines were estimated from 800 posterior draws and visualised as spaghetti plots. Thick black lines show median model-estimated proportion of antimicrobial-resistant E. coli across a range of observed temperature and humidity values, estimated from all 40000 posterior draws. For all splines the Y-axis shows log_10_ transformed estimated proportion of resistant isolates. Histograms above splines show the distribution of samples across the range of temperatures (left) or humidities (right). Note the differing y-axis range for cephalexin (AY and HS) and streptomycin (HS only)*.

For all 8 models, the ELPD for the models using a tensor-product smooth of thin-plate regression splines for temperature and humidity covariates was larger than for the models containing linear effects for these (Table S4). The increase in model performance suggests a non-linear relationship between these predictors and the outcome better describes the true relationship than a linear relationship.

### Temperature and humidity are positively associated with AMR

Across almost all antimicrobials and locations, there was evidence that temperature and humidity had a positive association with the proportion of resistant *E. coli* isolates within the farm environment samples. (Figure 4). Additionally, there was some evidence of a positive interaction effect between temperature and humidity, with warmer, more humid conditions associated with increased proportions of resistant isolates compared to cooler, drier conditions (Figure 4).

To demonstrate the effect of changing weather conditions, the model-estimated proportion of resistant isolates was calculated for 4 different weather condition change scenarios: increase in temperature, increase in humidity, positive interaction between temperature and humidity, and negative interaction between temperature and humidity (Table S5; Figure 5).

**Figure 5:**
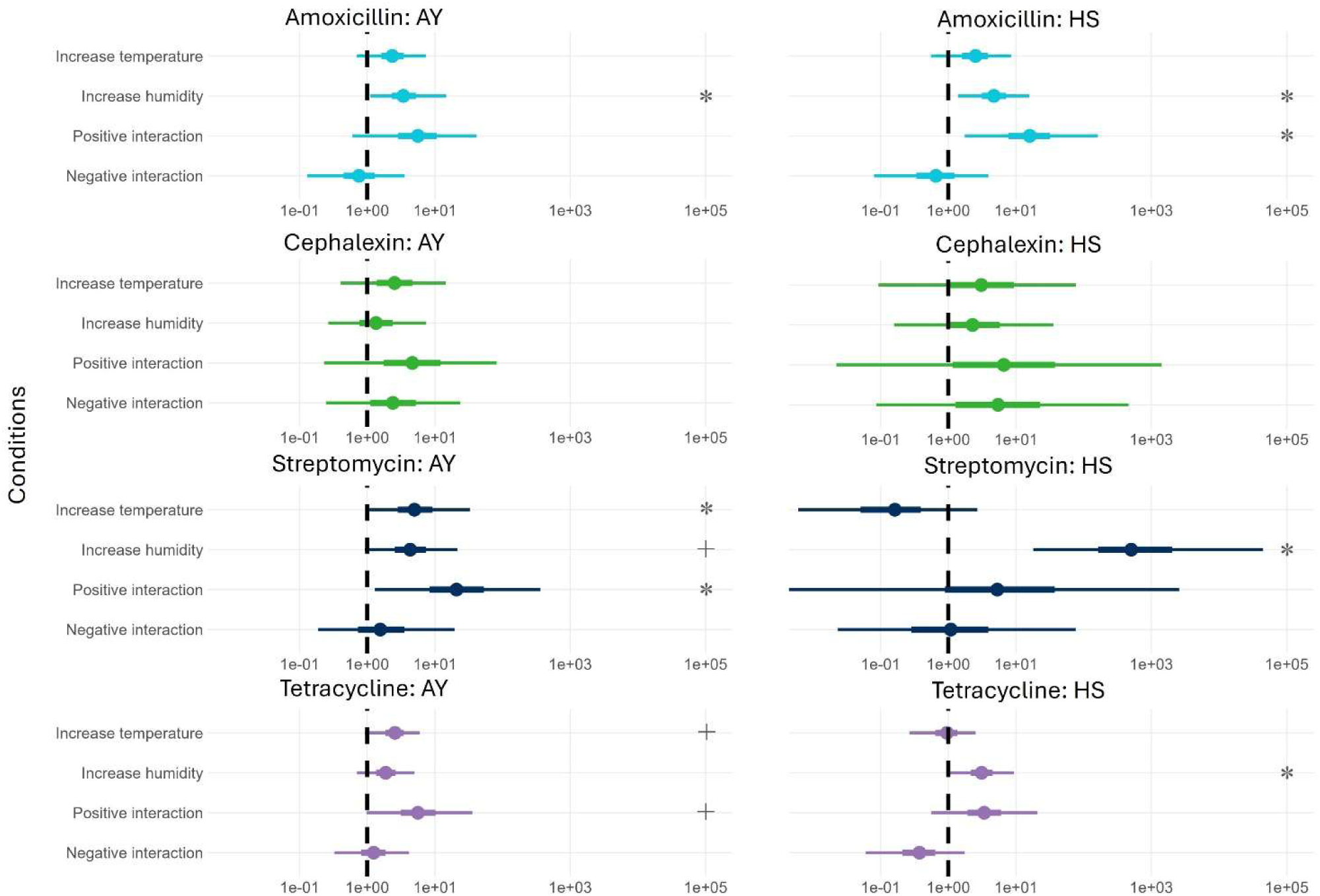
Change in model-estimated proportion of antimicrobial-resistant *E. coli* across different humidity and temperature values. *Points show median estimates, thick bars show 50% credible intervals, thin bars show 95% credible intervals. Proportion change calculated from the model-estimated proportion of antimicrobial-resistant E. coli across four weather condition change scenarios*. *1) ‘Increase temperature’: Increase from 1°C to 21°C, at 82% humidity* *2) ‘Increase humidity’: Increase from 60% to 92% humidity, at 10°C* *3) ‘Positive interaction’: From cold & dry (1°C, 60%) to hot & humid (21°C, 92%)* *4) ‘Negative interaction’: From cold & humid (1°C, 92%) to hot & dry (21°C, 60%)* *\* 95% credible interval does not cross the null of 1* *+ 90% credible interval does not cross the null of 1*

#### Streptomycin

Model estimates under different weather conditions provided strong evidence of an association between weather conditions and resistance to streptomycin. In the AY sample group, an increase in temperature from 1°C to 21°C was associated with a 5.0-fold increase in the proportion of streptomycin resistant isolates (median; 95% CI: 1.03 – 33.1; Table S5). Weaker evidence, where the 90% credible interval range excluded 1, although the 95% credible interval range spanned one, suggesting greater uncertainty surrounding the association was also observed for humidity. A 32% increase in humidity from 60% to 92% was associated with a 4.3-fold increase in the proportion of streptomycin resistant isolates (median; 90% CI: 1.2 – 16.4; 95% CI: 0.9 – 21.4). Additionally in the AY group, an increase in both temperature and humidity (from 1°C and 60%, to 21°C and 92%) was associated with a 20.9-fold increase in the proportion of streptomycin resistant isolates (median; 95% CI: 1.3 – 362.8), suggesting a synergistic effect between temperature and humidity.

In the HS sample group, humidity was strongly associated with resistance to streptomycin; an increase in humidity from 60% to 92% was associated with a 505-fold increase in the proportion of streptomycin-resistant isolates (median; 95% CI: 18.0 – 44,592.9). There was little evidence of an association with temperature or a synergistic interaction between temperature and humidity in the HS group.

#### Amoxicillin

Strong evidence of an association with humidity was also found for the HS sample group for amoxicillin resistance. The same increase in humidity (from 60% to 92%) was associated with a 4.7-fold increase in the proportion of amoxicillin resistant isolates (median; 95% CI: 1.4 – 15.7). An increase in both temperature and humidity (from 1°C and 60%, to 21°C and 92%) was associated with a 16.0-fold increase (median; 95% CI: 1.7 – 162.4). In the AY sample group, there was evidence of a positive association between humidity and resistance to amoxicillin (median: 3.4, 95% CI: 1.1 – 14.7).

#### Tetracycline

Here there was less certain evidence of an association between temperature and tetracycline resistance in the AY group; an increase in temperature (1°C to 21°C) was associated with a 2.6-fold increase in the proportion of tetracycline resistant isolates (median; 90% CI: 1.1 - 5.2; 95% CI: 0.9 – 6.0). Additionally, there was some evidence of a synergistic effect between temperature and humidity, as an increase in both (from 1°C and 60%, to 21°C and 92%) was associated with a of 5.6-fold increase (median; 90% CI: 1.3 – 25.5, 95% CI: 0.98 – 35.7). In the HS group, there was evidence of an association between humidity and proportions of antimicrobial-resistant *E. coli* colonies. Increasing humidity (from 60% to 92%) was associated with a 3.1-fold increase in the proportion of antimicrobial-resistant (95% CI: 1.1 – 9.4).

#### Cephalexin

In both sample groups, model estimates suggested that temperature and humidity were slightly positively associated with resistance to cephalexin. However, the effect size for these were small and the uncertainty around the estimates high, so overall evidence from this study for an association between cephalexin resistance with weather conditions is uncertain.

#### Differences exist between sample groups

The model estimates under different weather conditions suggest some differences between the 2 sample groups (AY and HS). In the AY group, higher temperatures were associated with increased proportions of resistant isolates for streptomycin and tetracycline, while no association between temperature and the proportion of resistant *E. coli* was observed for any antimicrobial in the HS group. In contrast, the influence of humidity on the proportion of resistant isolates appeared greater in the HS group than in the AY group.

#### Some seasonal variation is unexplained by temperature and humidity

The spline included for ‘day of study’ revealed that there was some residual variation in resistance across the study which was not accounted for by changes in local weather (Figure S10). In the cephalexin models, a notable pattern of increases, and decreases in the proportion of antimicrobial-resistant *E. coli* across both study years could be observed. This suggests that some unmeasured seasonal factor may be further influencing the proportion of antimicrobial-resistant *E. coli* in these samples. This cyclic seasonal variation was not seen in any other antimicrobials when non-linear effects for temperature, humidity and their interaction were included in the models, although variability in the proportion of antimicrobial-resistant *E. coli* across the study period was observed.

## Discussion

This study found evidence that local temperature and humidity was associated with the proportion of *E. coli* isolates resistant to amoxicillin, streptomycin and tetracycline found in faecal samples from adult dairy cattle collecting yards and heifers shed housing. This is the first study the authors are aware of that assesses relative humidity as a predictor for livestock AMR in addition to temperature. The analysis found evidence that humidity was positively associated with resistance, particularly in the HS (heifers in sheds/pens) sample group. This could potentially be due to higher humidity preventing the desiccation of environmental *E. coli,* increasing survival and likelihood of AMR acquisition. Additionally, there was some evidence for a synergistic effect between temperature and humidity with warmer, damper conditions being associated with higher proportion of resistant isolates.

Our study found evidence for large relative changes in the proportion of AMR with temperature and humidity. Although the likelihood of sample-level positivity was quite high, the proportion of antimicrobial-resistant *E. coli* within the farm environment samples was generally low. This resulted in small absolute changes in the proportion of resistant *E. coli*, which are arguably unlikely to cause significant increases in AMR infections and antimicrobial treatment failures in the cattle populating these environments. However, the prevalence of AMR bacteria on ruminant farms in the UK and globally is unclear as no representative surveillance has been carried out on farms directly. In environments with a higher baseline prevalence of AMR bacteria, even a few-fold increase in resistance could pose a significant risk to the health of humans and animals, especially considering the predicted increase in global temperatures due to climate change (34).

The observational study design does not provide evidence of there being a causal relationship between local weather conditions preceding sampling and AMR prevalence. Other factors which potentially contribute to the correlations observed could include seasonal changes in disease burden, antimicrobial usage or disease outbreaks (e.g. heat stress in summer, summer mastitis due to flies, respiratory infections in winter) and farm management practices.

An alternative explanation for the association with humidity in the HS group, for example, could be farm management activities, which were unmeasured in this study. During periods of higher rainfall, and therefore higher relative humidity due to evaporation, grazed cattle might be brought into indoor housing to prevent poaching of the grazing land. This could increase the stocking density of indoor housing, leading to higher faecal pollution of their environment and warmer, more humid conditions inside cattle housing, in turn potentially increasing the proportion of antimicrobial-resistant *E. coli*. Movement of cattle on the farm and stocking density of the sampled areas was not captured within this study, so this could explain some of the association seen between humidity and AMR.

Additionally, these factors could lead to an increased abundance of *E. coli,* increased antimicrobial use at these times, or a perturbed microbiome (35), allowing antimicrobial-resistant bacteria which would otherwise be subject to fitness costs to colonise the environment at higher rates. Another possible explanation for the relationship between elevated temperature and higher prevalence of resistance might be increased water consumption of cattle. Water troughs have previously been identified as on-farm reservoirs for pathogenic *E. coli* as well as antimicrobial-resistant *E. coli*, so, at times of increased water consumption by cattle, the abundance of *E. coli* in the gut and the faecally contaminated environment might also increase (19, 36). Dilution of *E. coli* concentrations resulting from increased water consumption and provision under warmer conditions may however also occur and so further research to elucidate the dynamics of *E. coli* abundance and proportions of antimicrobial-resistant *E. coli* at the water trough interface may be warranted.

Outdoor air temperature and humidity data used in this study were measured at the nearest weather station and therefore and may not accurately reflect conditions at the sampling sites of the collecting yard and indoor cattle housing. The magnitude of potential measurement error might have varied between farms and sample locations. This may have led to higher uncertainty being observed in the coefficient estimates within the indoor housing (HS) sample group models where, the temperature and humidity were likely to have differed from outdoor sampling sites.

Seven-day means were calculated for the weather predictors from hourly observations. Previous studies have shown that *E. coli* can survive in cattle manure and soil for several weeks (37, 38, 39, 20). As the farm sample sites were likely exposed to continual faecal contamination over time, the microbiome of the farm environment could be shifting over weeks or months and possibly be modulating AMR. Therefore, it is possible that AMR is more greatly impacted by longer-term weather trends, such as those over multiple weeks or months, rather than the over the 7-day period considered. Testing additional time periods and applying a variable selection technique could reveal stronger associations between the weather predictors and AMR.

In addition to potential measurement error in the weather data, measurement error may also exist in the colony counts produced by the microbiology method. This is particularly true for plates with high colony counts which were more likely to have overlapping colonies or which were estimated from a count of a subsection of the plate (for plates >300 colonies). Colony counting error, however, has a lower relative impact on the estimated proportion of resistant isolates in samples with high *E. coli* abundance (and therefore high colony counts) than those with low abundance. Therefore, the error introduced due to miscounting colonies should have had limited impact on the overall findings.

Finally, it should be considered that the associations observed between local weather conditions and AMR could be due to the effect of temperature on bacterial growth and prevalence. The microbiology methodology and Poisson likelihood modelling approach used in this study aim to control for the effect of bacterial growth on the likelihood of detecting resistance. Despite the utilisation of bacterial abundance within the model, it remains that there is a relative paucity of information in samples with low *E. coli* abundance in comparison to those with high abundance, and therefore the likelihood of detecting any resistance remains higher in high-abundance samples.

To further investigate the effect of weather on AMR in the farm environment, future studies could employ a more detailed design in which weather parameters such as temperature, humidity, rainfall and ultraviolet light exposure (via sunlight) are monitored continuously at the sample site. If such a design was employed, recording any antimicrobial treatments, disease prevalence and the number and density of cattle populating the sample site would allow these potential confounding factors to be controlled for within statistical analysis.

Caution is suggested in extrapolating these results to other farm systems, other climates or other bacterial species. Only dairy farms were included in our study, and these were a convenience sample whose participation in the project was voluntary. Willingness to agree to participation may be associated with the farmers who are both more aware of AMR and who believe they use antimicrobials in a responsible way, as the wider study involved farm management and antimicrobial use data collection (19). It is therefore difficult to generalise our results beyond dairy farms with similar management practices and climate to those of South-West England. Finally, only *E. coli* were cultured and tested for AMR. The effect of weather on resistance *E. coli* could differ from that of other bacteria which occupy different niches or have different physiology or resistance mechanisms, such as Gram-positive bacteria.

This study provides important insights into AMR prevalence throughout the year and in changing environmental conditions. The findings add to the increasing body of evidence that temperature and AMR are associated, such as those found in humans in the US and Europe (10, 12). This has implications for the combined global health concerns of climate change and AMR.

Understanding the interplay between environmental factors and AMR is beneficial in unpicking the risk factors which drive AMR, and thereby the interventions which have the potential to reduce the risk of AMR development. In addition, understanding the association between weather and seasonality on AMR is essential for designing effective risk factor studies or surveillance programs within agricultural environments. The reason for this is 2-fold. The evidence presented in this study suggests that samples taken in periods with warmer, more humid weather conditions contain a higher proportion of antimicrobial-resistant *E. coli* on average. As well as directly influencing AMR, the effect of weather conditions on *E. coli* abundance could impact the likelihood of detecting resistance if samples are not standardised or the results not analysed in a way which accounts for differing *E. coli* abundance. Commonly used microbiology techniques which are insensitive to the proportion of resistant bacteria and only detect presence or absence will be especially impacted by the differing *E. coli* abundance. An improved understanding of the complex relationships between environmental factors and AMR is essential for designing surveillance systems and estimating the risk AMR poses to animal health, human health and food security in the future.

## Acknowledgements

We thank all the farmers and veterinary surgeons who participated in this study. In addition, we thank the milk processing companies who supported our recruitment process and the veterinary students who helped with data processing and sample collection.

The One Health Selection and Transmission of Antimicrobial Resistance project was funded by grant NE/N01961X/1 from the Antimicrobial Resistance Cross Council Initiative supported by the seven United Kingdom Research Councils. Elliot Stanton was supported by the Biotechnology and Biological Sciences Research Council-funded South West Biosciences Doctoral Training Partnership [training grant reference BB/T008741/1].

## Supplementary

**Figure S1:**
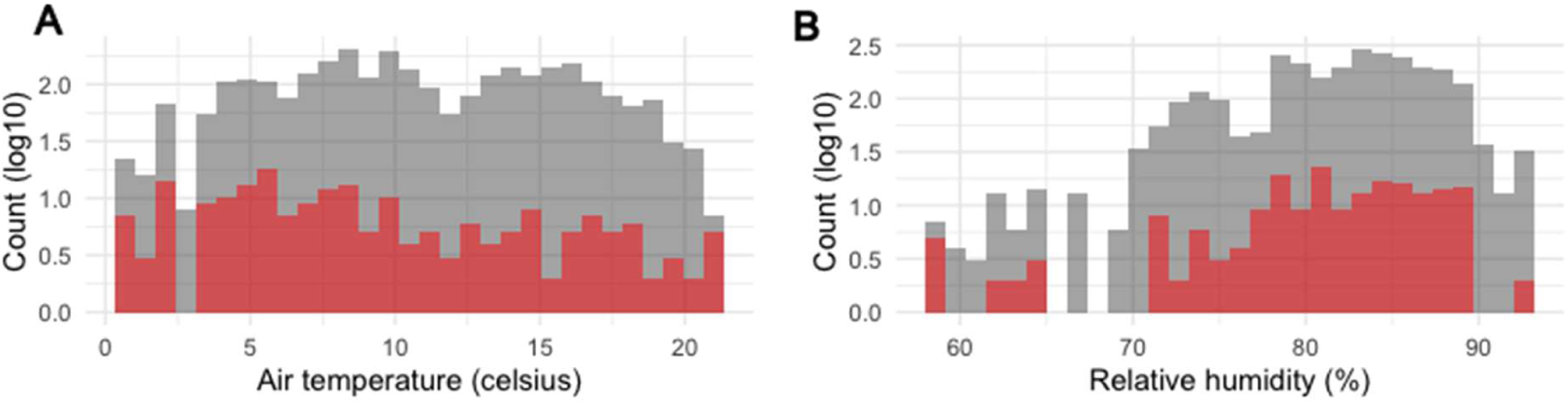
Histograms showing the distribution of samples with no *E. coli* cultured across the range of weather conditions. *Red: samples excluded from analysis due to no E. coli cultured. Grey: samples included in the analysis. Y-axis is log_10_ plus 1 transformed. Temperature and relative humidity reflect the mean across hourly observations in the 7-day period preceding sample collection.*

**Figure S2:**
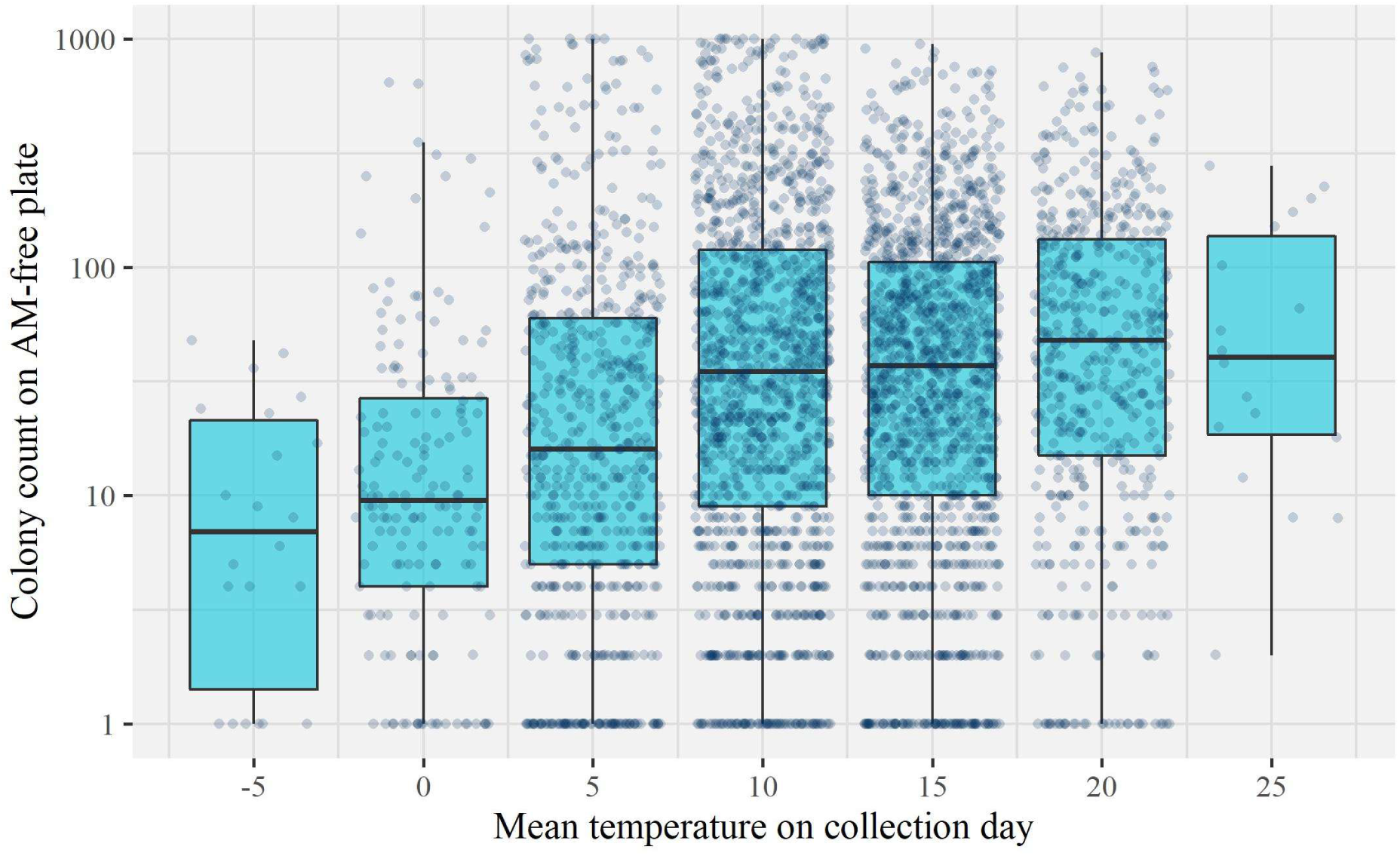
Correlation between E. coli abundance and mean temperature (°C) on day of sample collection. *Box plots to show the distribution of CFU on the AM-free plate for each sample grouped into 5°C intervals of the mean temperature on the day of collection. Dark points show the CFU of individual samples. Y-axis is log_10_ transformed. Colony counts of zero are not shown.*

**Figure S3:**
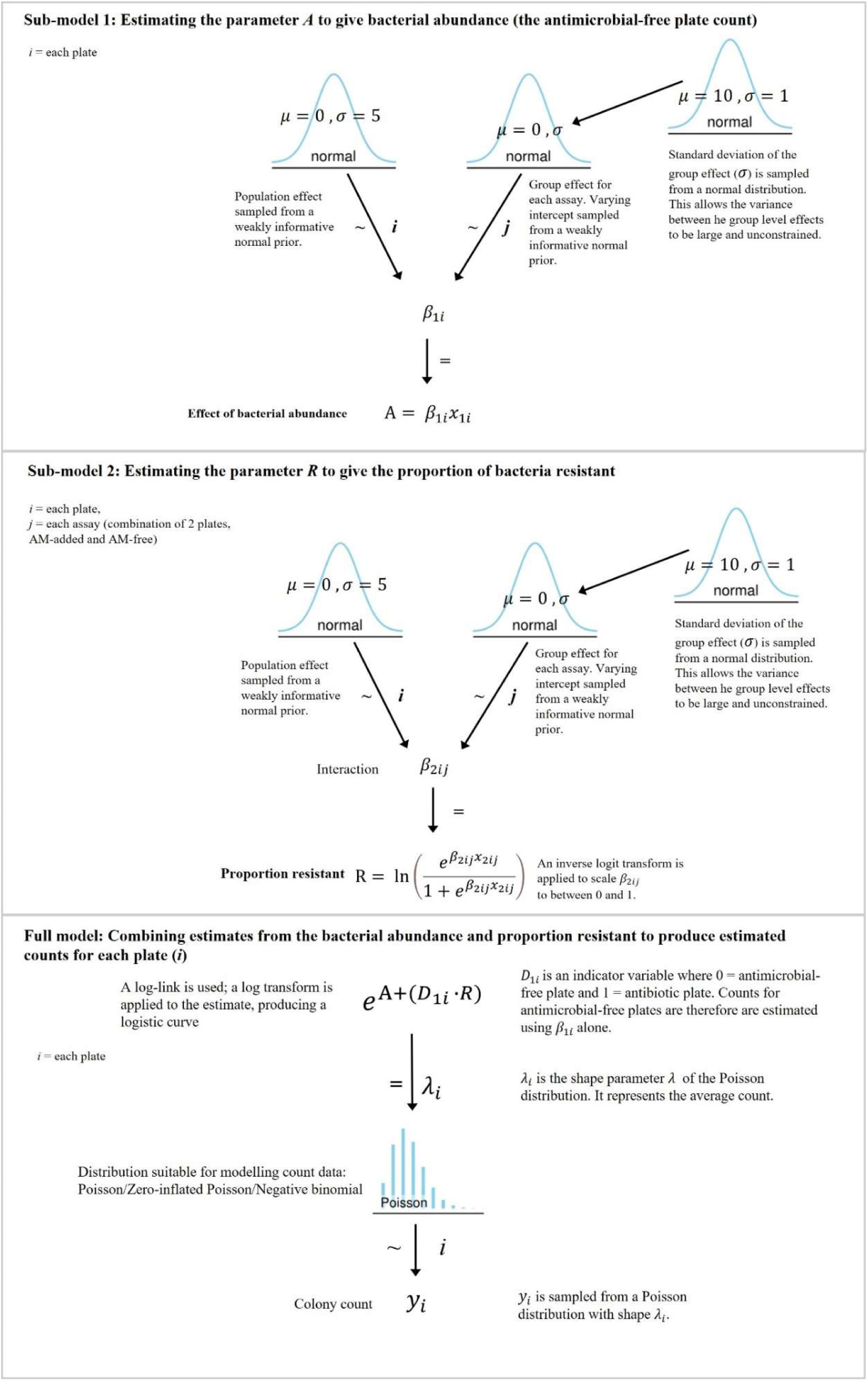
Schematic showing model structure. *Adapted from (1) Sub-model 1 provides estimates the overall abundance of E. coli for a given combination of AM-free and AM added agar plates (A). Sub-model 2 estimates the proportion of antimicrobial-resistant E. coli against the antimicrobial contained within the AM added agar (R). An inverse logit transform is used to scale estimated proportions between 0 and 1 under the assumption that the number of antimicrobial-resistant E. coli colonies cannot exceed the total E. coli abundance. Estimates of A and R are combined within the full model to estimate a count for each agar plate.*

**Table S1:** Prior choice for non-linear Bayesian model with Poisson likelihood for the proportion of AMR isolates.

| Parameter<br>name | Parameter<br>description | Effect | Prior distribution |
| --- | --- | --- | --- |
| interaction | Nonlinear parameter | Population effect | Normal($\mu = 0, \sigma = 5$) |
| | for proportion of | Group effect | Normal($\mu = 10, \sigma = 1$) |
|  | resistant bacteria | standard deviation |  |
| main | Nonlinear parameter | Population effect | Normal($\mu = 0, \sigma = 5$) |
| | for bacterial abundance | Group effect | Normal($\mu = 10, \sigma = 1$) |
|  |  | standard deviation |  |

**Figure S4:**
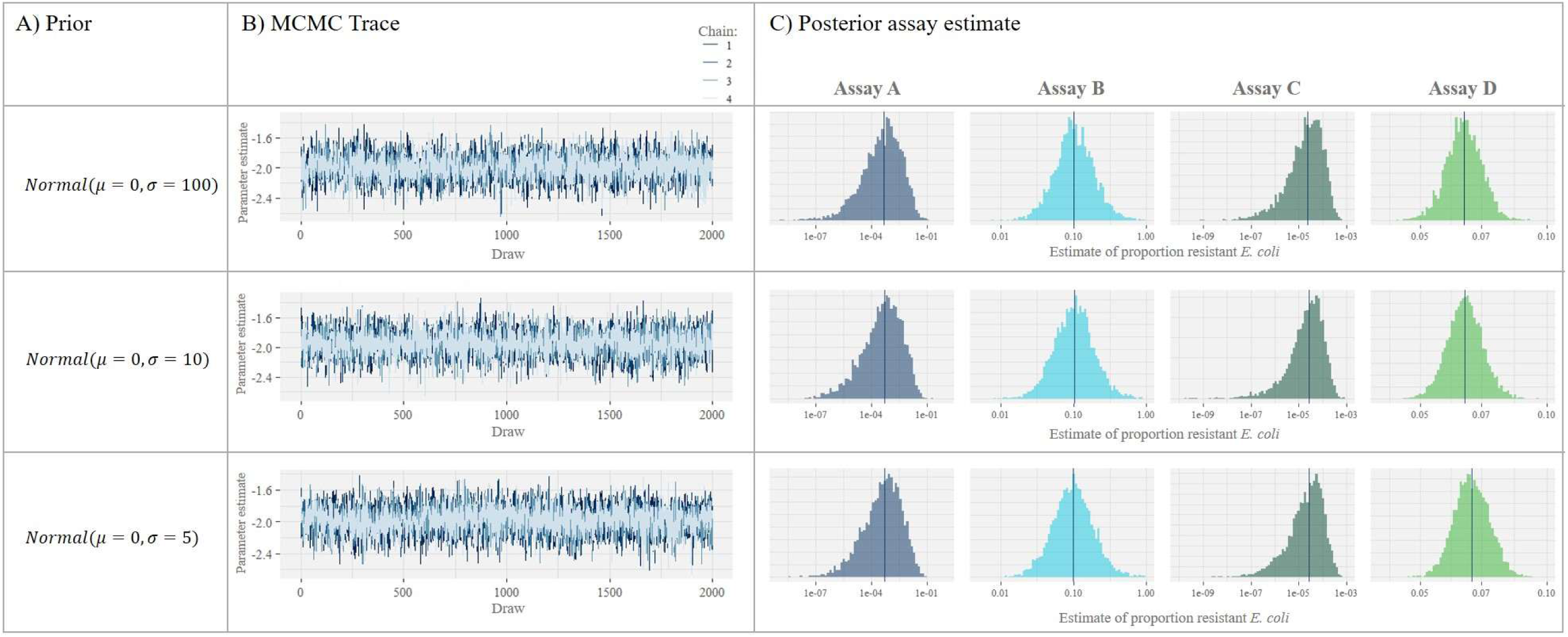
Prior sensitivity analysis. *A) Prior distribution tested. B) Untransformed parameter estimates for a randomly chosen assay. All 2000 draws (inclusive of warm-up) from each of the four MCMC chains are shown. C) Distribution of parameter estimates for four randomly chosen assays shown on the transformed scale – the proportion of resistant E. coli. The vertical blue line shows the mean estimate. Assays with a range of counts were selected: Assay A – low AM-free count, zero AM count. Assay B – low AM-free count, >0 AM count. Assay C – high AM-free count, zero AM count. Assay D – high AM-free count, >0 AM count.* *Prior distributions for the non-linear parameter, ‘main’, estimated in sub-model 1 and ‘interaction’, estimated in sub-model 2 were chosen to be weakly informative in line with current ‘best practice’ in Bayesian statistics and to aid model compilation (2, 3). The inverse logit transform applied to the ‘interaction’ parameter limited the result to between 0 and 1. On the inverse logit scale, very negative estimates tend towards zero and high positive estimates tend toward one. Choosing a very uninformative prior would therefore allow explorations of very high or low estimates which would provide little extra information to the model. As such, normal priors with limited standard deviations were tested. To ensure the prior was not unexpectedly biasing the results, a variety of normal priors centred about 0 with different standard deviations were trialled; normal(μ=0,σ=100), normal(μ=0,σ=10), normal(μ=0,σ=5) (A). No irregularities such as poor mixing or incomplete sample space exploration were seen. The convergence diagnostic, the potential scale reduction factor (R̅), was <1.05 for all parameters in each of the 3 models (B), suggesting that HMC chains had appropriately sampled the solution space (4, 5). The distribution of estimates for the proportion of antimicrobial-resistant E. coli in each model were consistent, suggesting the most informative prior, normal(μ=0,σ=5), was not biasing the effect size in any direction or reducing the variance of the estimate.*

**Figure S5:**
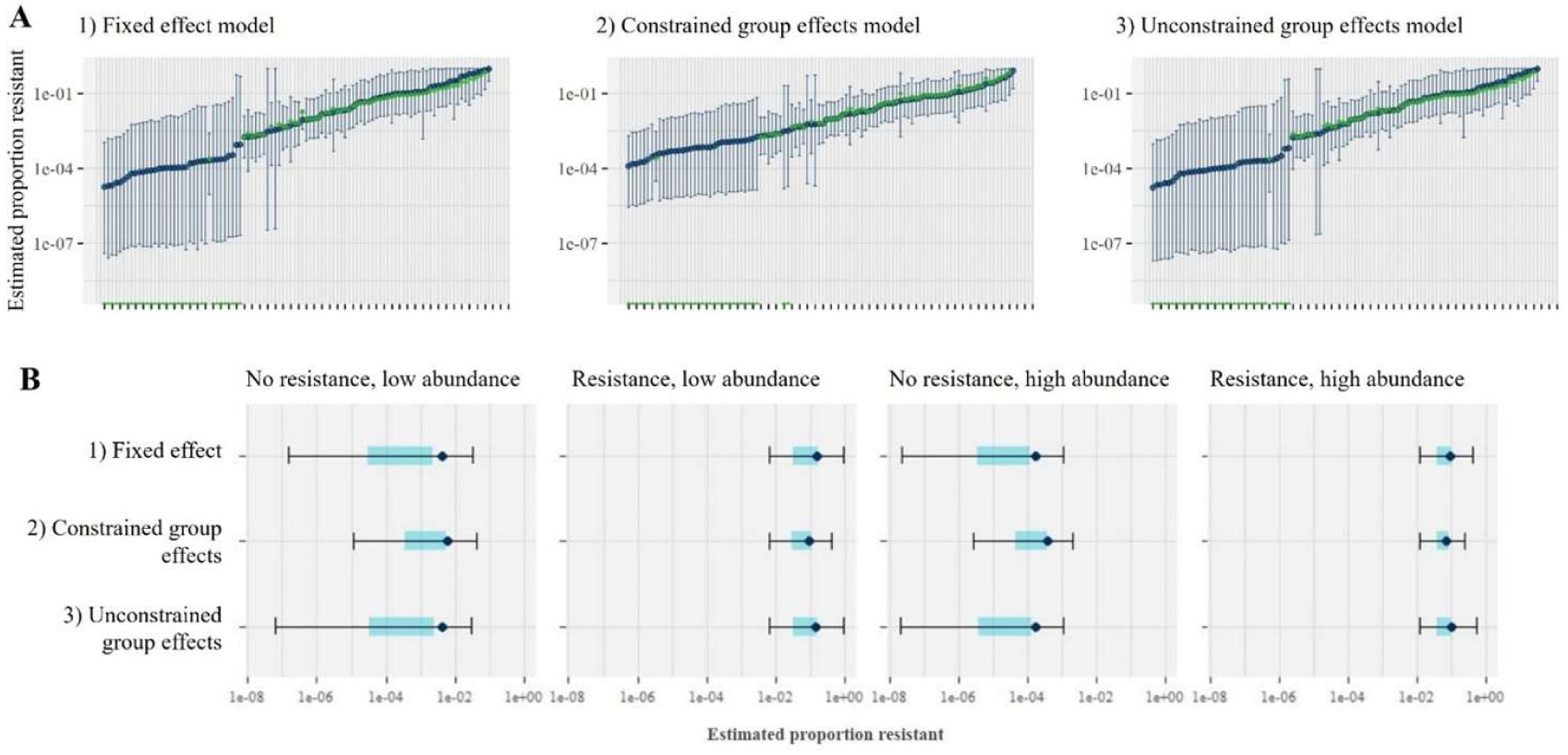
Comparing fixed effect to group effects with varied priors. A) Posterior predictive checks for a subset of 100 randomly chosen samples from each model. Blue points show the mean estimate of the proportion antimicrobial-resistant E. coli for each assay with 95% credible intervals as error bars. Green points are the proportion of antimicrobial-resistant E. coli calculated from the raw data. Assays ordered left to right by the mean estimate of proportion resistant. B) Boxplots showing the distribution of draws for four randomly chosen assays with a range of bacterial abundances and AMR levels across the three models. Blue points show mean, the shaded region shows IQR and whiskers show 95% CI. Model 1) Fixed effect for assay intercept. This allows intercept to vary between assays and is estimated separately for each assay. Model 2) Group effect for assay intercept with a half T Student prior for the standard deviation between assays, constraining the amount of variation and assuming the assay effects are normally distributed. Model 3) Group effect for assay intercept with a normal(10,1) prior for the standard deviation of the intercepts, effectively unconstraining the amount of variation between samples. Including a group effect for each assay (a pair of AM-free and AM-added counts observed for a sample) allowed the intercept to vary between each pair of counts. Initially this group effect was sampled from a normal prior centred around zero, constraining variation between samples to be normally distributed with low variance. This pattern of variance was not obvious within the data and so to lessen this assumption but retain the group effect structure, the priors determining the variance of the assay group effect were changed to allow greater variation between samples. The standard deviation of the assay group effect was widened to be weakly informative by changing its prior from a restrictive half T-Student distribution to a normal distribution with a mean of 10. Comparison of the original prior (constrained group effects), the updated prior (unconstrained group effects) and a model which parameterised assay as a fixed effect, which therefore had no constraint on variation between assays, indicated that the constrained group effect had smaller 95% CI ranges compared to a fixed-effect model. In contrast, 95% CI ranges within a model using the unconstrained group effect priors produced similar estimated ranges as the fixed-effect parameterization, indicating that this prior was able to unconstrain the variation between assays. This effect was observed across a range of assays with differing E. coli abundance and proportions of antimicrobial-resistant E. coli.

**Figure S6:**
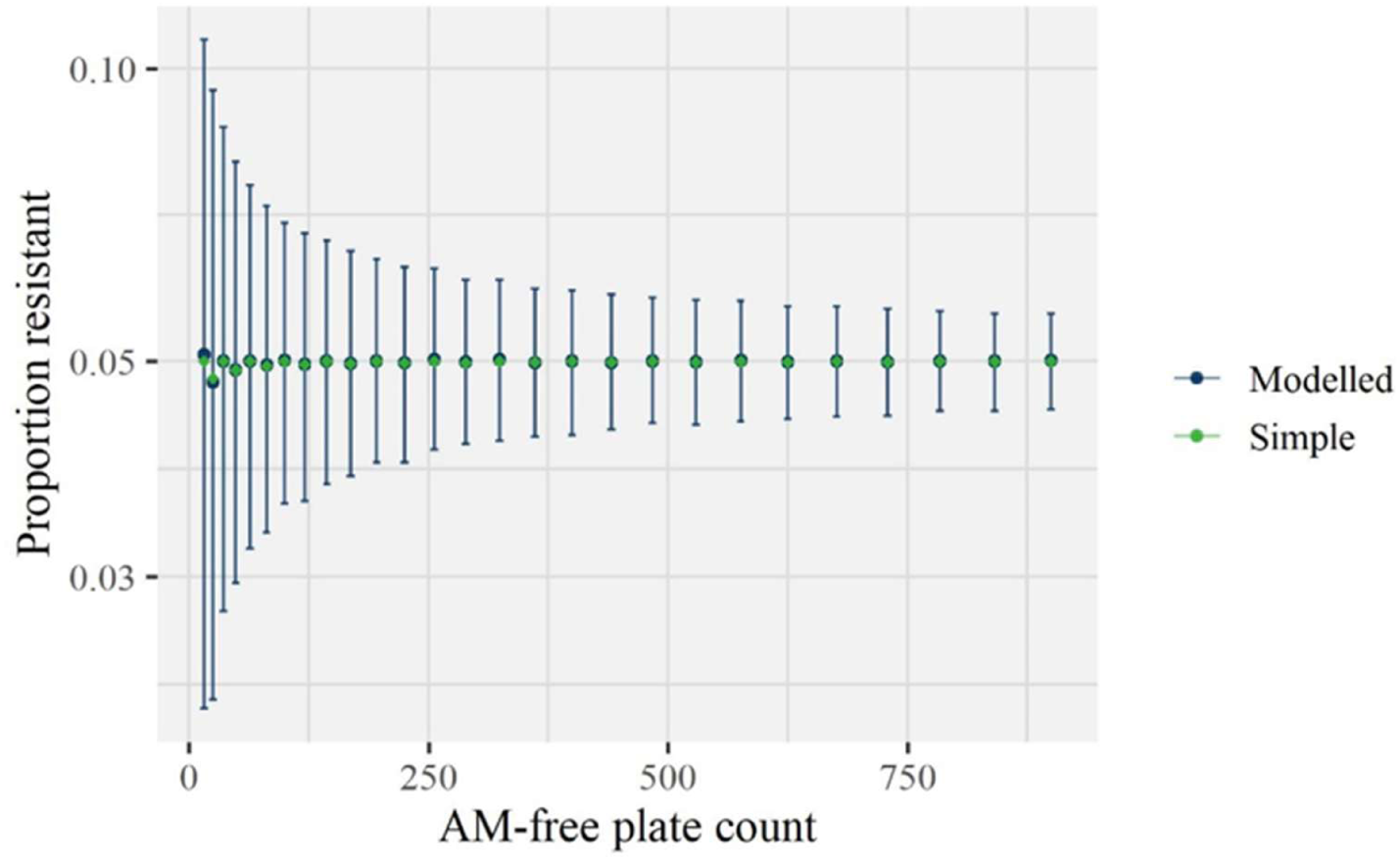
Uncertainty around the estimate of proportion antimicrobial-resistant E. coli decreases as E. coli abundance increases when modelling a synthetic dataset. A subset of results from modelling of synthetic data where the estimated proportion resistant ≈0.05. Error bars show 95% credible interval, and points show the mean estimate. Light points show the proportion resistant as calculated from the raw data. Y-axis is log base 10 transformed. Uncertainty around the proportion of antimicrobial-resistant E. coli within samples was greatest when E. coli abundance was lower. As E. coli abundance increased the uncertainty of estimates of the proportion of antimicrobial-resistant E. coli decreased. The mean model estimates closely match the proportion of antimicrobial-resistant E. coli estimated by dividing the AM-added colony count by the AM-free count, with 95% CI’s including this proportion at all AM-free counts.

**Figure S7:**
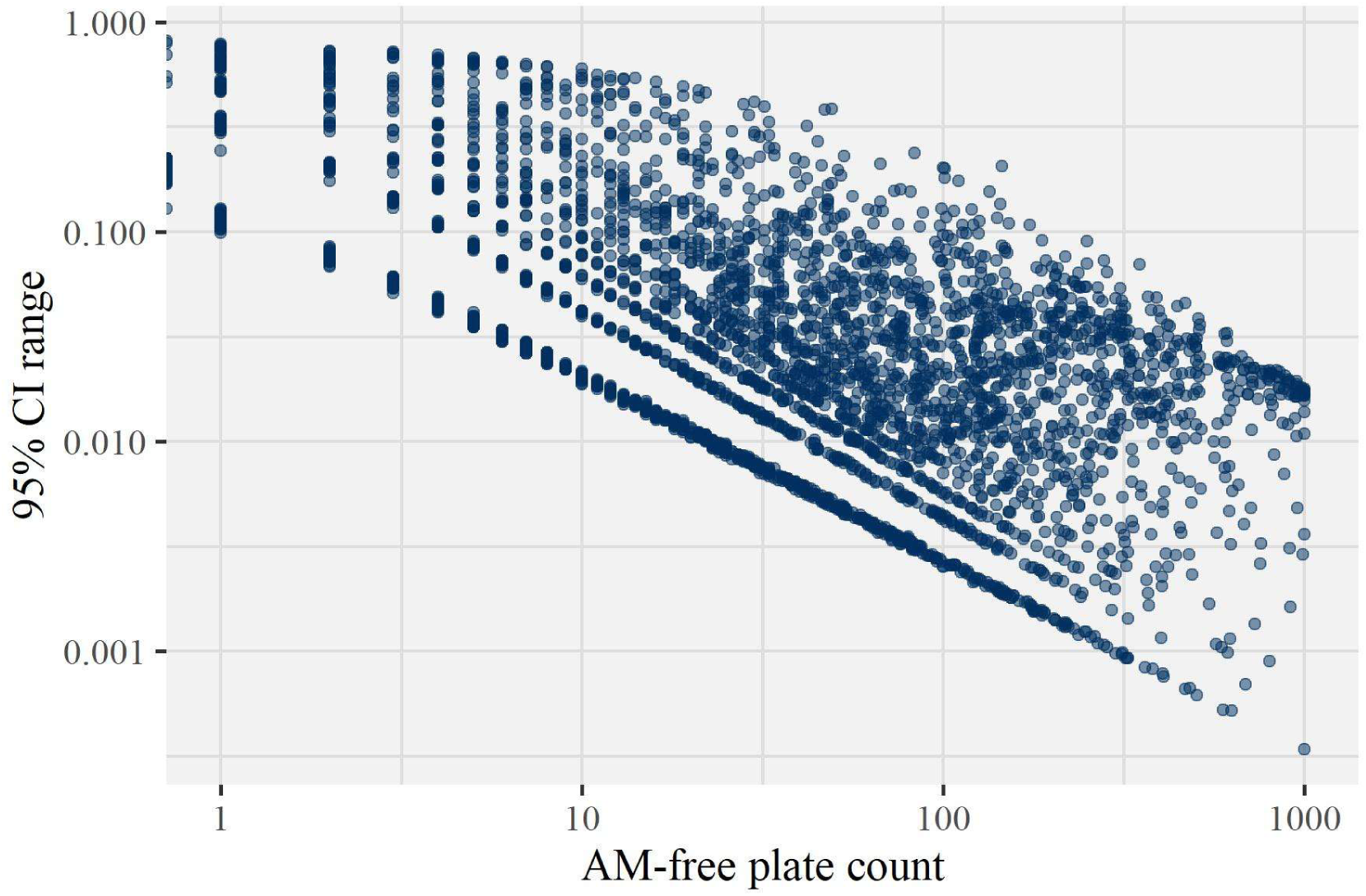
Uncertainty around the estimate of proportion of antimicrobial-resistant E. coli decreases as E. coli abundance increases when modelling the proportion of tetracycline-resistant E. coli. The 95% credible interval range for the modelled proportion of resistant E. coli was calculated and is plotted against the raw colony count on the AM-free plate. Banding on the x and y axes is due to the discrete count data for AM-free and AM-added counts; multiple samples had the same plate counts. Both axes are log_10_ transformed.

**Figure S8:**
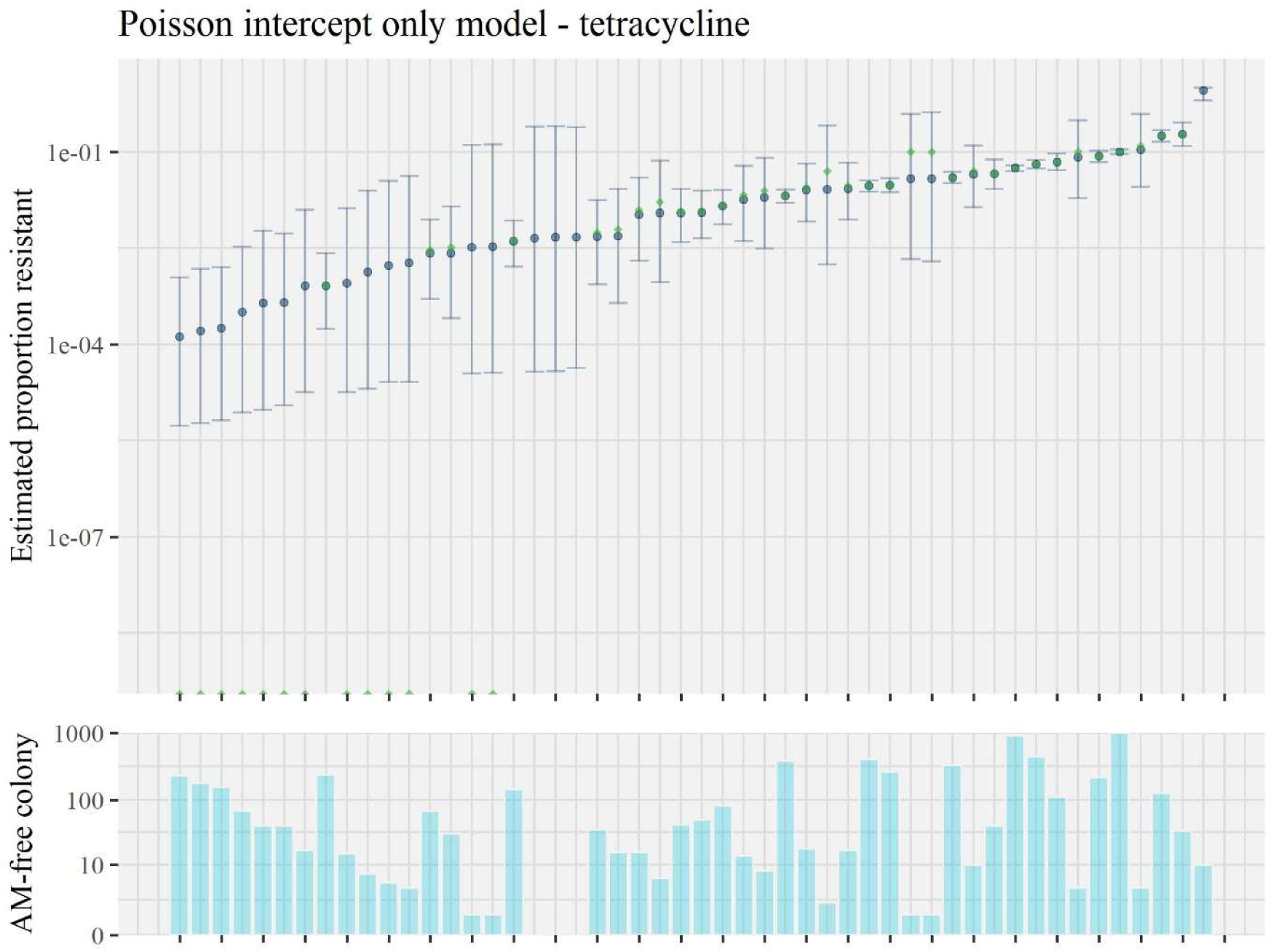
Posterior predictive check of the intercept-only model for tetracycline resistance. Estimates for the proportion of tetracycline-resistant E. coli in a random subset of 50 samples from the dataset. Blue points in the top subplot show the mean estimate of the proportion of antimicrobial-resistant E. coli for each sample from the model, with 95% credible intervals as error bars. Green points are the proportion of antimicrobial-resistant E. coli calculated through dividing the number of E. coli colonies observed to grow on AM-containing agar by those growing on AM-free agar. Green points on the x-axis equal zero and are from samples with no tetracycline-resistant colonies. Samples are ordered left to right by the mean estimated proportion of antimicrobial-resistant E. coli. Y-axis is log_10_ plus one transformed. The bottom subplot shows the raw colony count data from the AM-free plate for each assay, ordered the same as the plot above. Y-axis is log_10_ plus one transformed. The same relationship between AM-free colony counts and estimate uncertainty observed in Supplementary Figures 6 and 7 is observed here. Where no antimicrobial-resistant E. coli colonies are observed within a sample, estimated proportions of antimicrobial-resistant E. coli are low and uncertainty is high. In such cases 95% CIs extend beyond the hypothetical lower limit of detection of 1×10^-4^ (which would represent 1 E. coli colony being observed on an AM-containing plate while a count of 1000 was recorded for the corresponding AM-free agar-plate). Where no E. coli colonies are observed on either the AM-free or AM-added plates 95% CIs are very wide.

**Figure S9:**
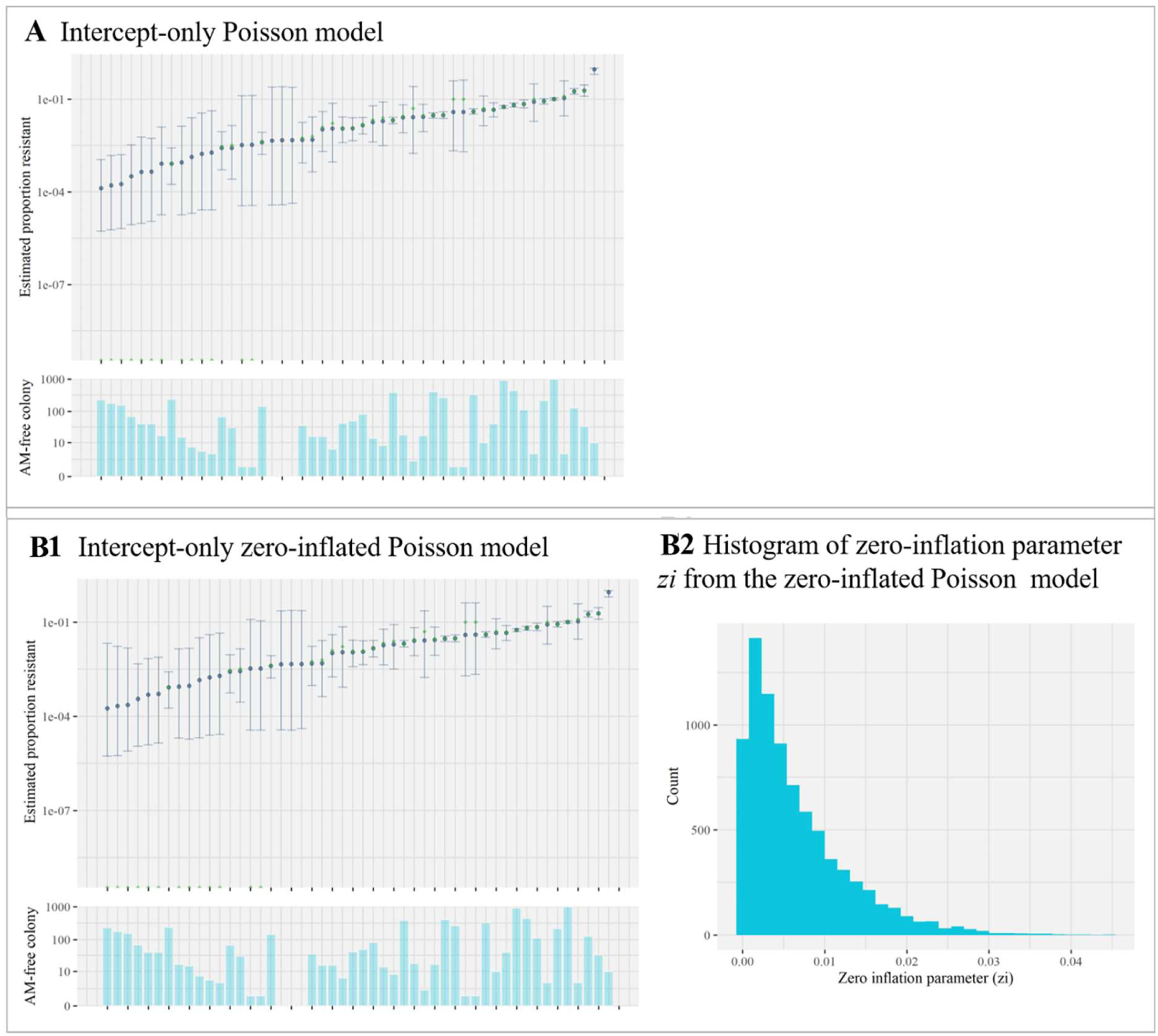
Comparison of two models for the proportion of antimicrobial-resistant E. coli with differing underlying distributions for colony count. A, B1) Models estimating the proportion of streptomycin-resistant E. coli in a subset of 500 samples. 50 randomly chosen samples are shown. In the top subplot, blue points show the mean estimate of the proportion of antimicrobial-resistant E. coli for each sample from the model with 95% credible intervals as error bars. Green points are the proportion of antimicrobial-resistant E. coli calculated through dividing the number of E. coli colonies observed to grow on AM-containing agar by those growing on AM-free agar. Green points on the x-axis equal zero and are from samples with no antimicrobial-resistant colonies. Samples are ordered left to right by the mean estimated proportion resistant. Y-axis is log_10_ plus one transformed. The bottom subplots show the raw colony count data from the AM-free plate for the corresponding assay result shown in the subplot above. Y-axis is log_10_ plus one transformed. B2) Histogram showing posterior draws for the zero-inflation parameter zi. Zi is scaled between 0 and 1, giving the proportion of zeroes model as excess.

**Table S2:**
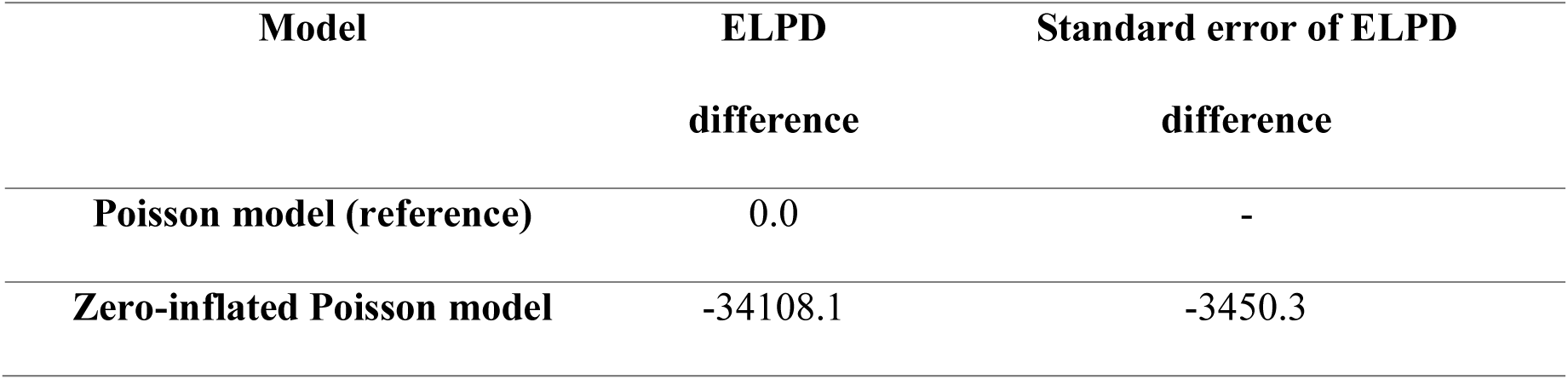
K-fold cross-validation. Stratified 20-fold cross-validation was run on the three models shown in Figure 8. The Poisson model had the highest expected log predictive density (ELPD) and is used as a reference for the comparison. The standard error of these differences was also calculated.

**Table S3:**
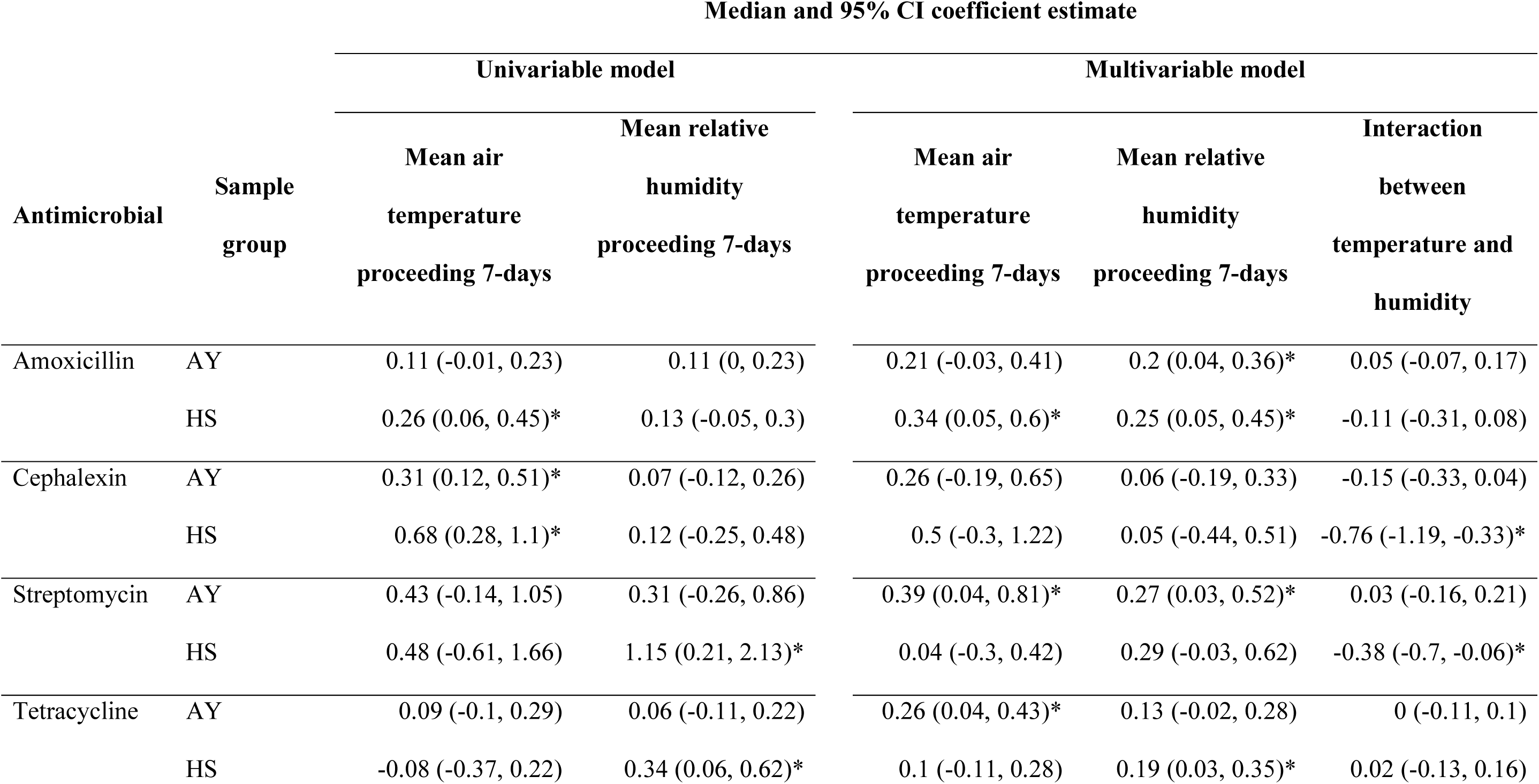
Coefficient estimates from both univariable and multivariable linear regression models for the effect of temperature and humidity on the proportion of antimicrobial-resistant *E. coli* in the farm environment samples. *Coefficient estimates are for the standardised predictors so give the effect on the proportion of antimicrobial-resistant E. coli when there is a 1 standardised unit increase in the predictor*. *\* Indicates when the 95% credible interval of the posterior doesn’t cross the null hypothesis (zero)*

| Antimicrobial | Sample group | Univariable model |  | Multivariable model |  |  |
| --- | --- | --- | --- | --- | --- | --- |
|  |  | Mean air temperature proceeding 7-days | Mean relative humidity proceeding 7-days | Mean air temperature proceeding 7-days | Mean relative humidity proceeding 7-days | Interaction between temperature and humidity |
| Amoxicillin | AY | 0.11 (-0.01, 0.23) | 0.11 (0, 0.23) | 0.21 (-0.03, 0.41) | 0.2 (0.04, 0.36)* | 0.05 (-0.07, 0.17) |
|  | HS | 0.26 (0.06, 0.45)* | 0.13 (-0.05, 0.3) | 0.34 (0.05, 0.6)* | 0.25 (0.05, 0.45)* | -0.11 (-0.31, 0.08) |
| Cephalexin | AY | 0.31 (0.12, 0.51)* | 0.07 (-0.12, 0.26) | 0.26 (-0.19, 0.65) | 0.06 (-0.19, 0.33) | -0.15 (-0.33, 0.04) |
|  | HS | 0.68 (0.28, 1.1)* | 0.12 (-0.25, 0.48) | 0.5 (-0.3, 1.22) | 0.05 (-0.44, 0.51) | -0.76 (-1.19, -0.33)* |
| Streptomycin | AY | 0.43 (-0.14, 1.05) | 0.31 (-0.26, 0.86) | 0.39 (0.04, 0.81)* | 0.27 (0.03, 0.52)* | 0.03 (-0.16, 0.21) |
|  | HS | 0.48 (-0.61, 1.66) | 1.15 (0.21, 2.13)* | 0.04 (-0.3, 0.42) | 0.29 (-0.03, 0.62) | -0.38 (-0.7, -0.06)* |
| Tetracycline | AY | 0.09 (-0.1, 0.29) | 0.06 (-0.11, 0.22) | 0.26 (0.04, 0.43)* | 0.13 (-0.02, 0.28) | 0 (-0.11, 0.1) |
|  | HS | -0.08 (-0.37, 0.22) | 0.34 (0.06, 0.62)* | 0.1 (-0.11, 0.28) | 0.19 (0.03, 0.35)* | 0.02 (-0.13, 0.16) |

**Table S4:**
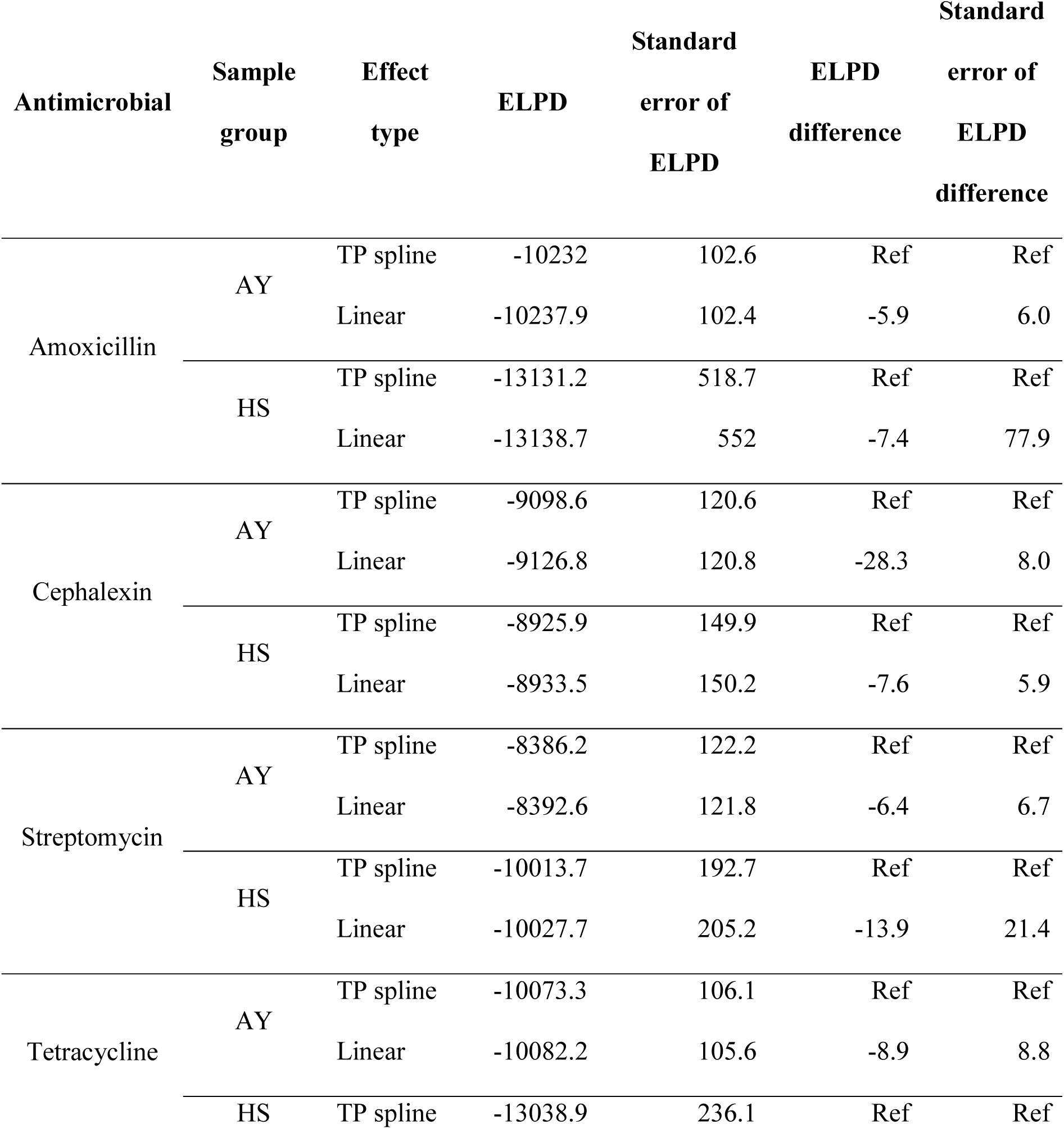
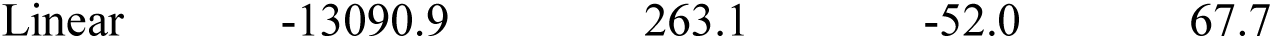
Expected log predictive density of models with linear effects versus thin-plate regression splines for weather variables. *AY: Adults in collecting yard* *HS: Heifers in sheds/pens* *ELPD calculated from k-fold cross validation using 5 folds. Where ELPD is larger between two models with the same data and structure, the model has greater predictive performance*.

**Table S5:**
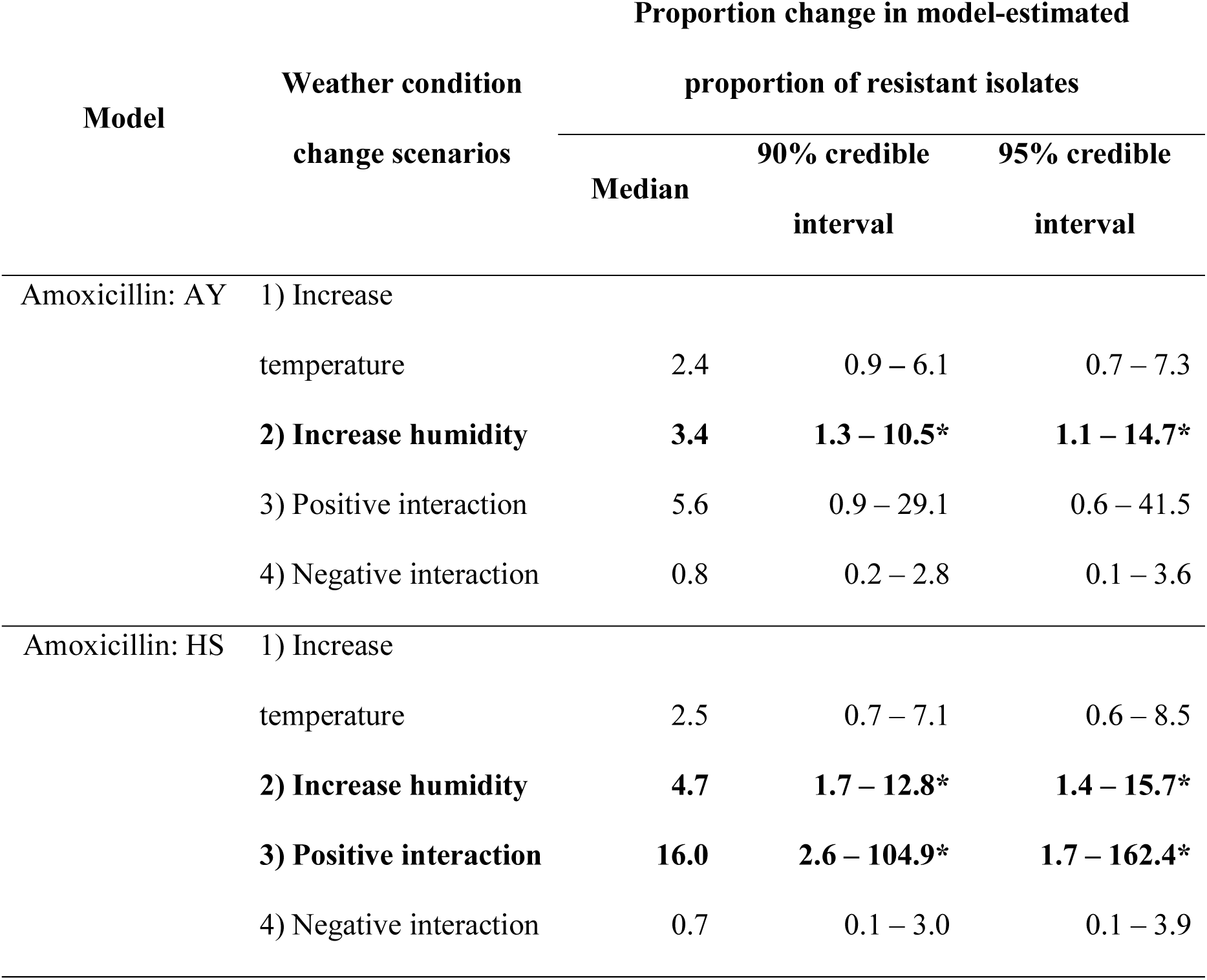
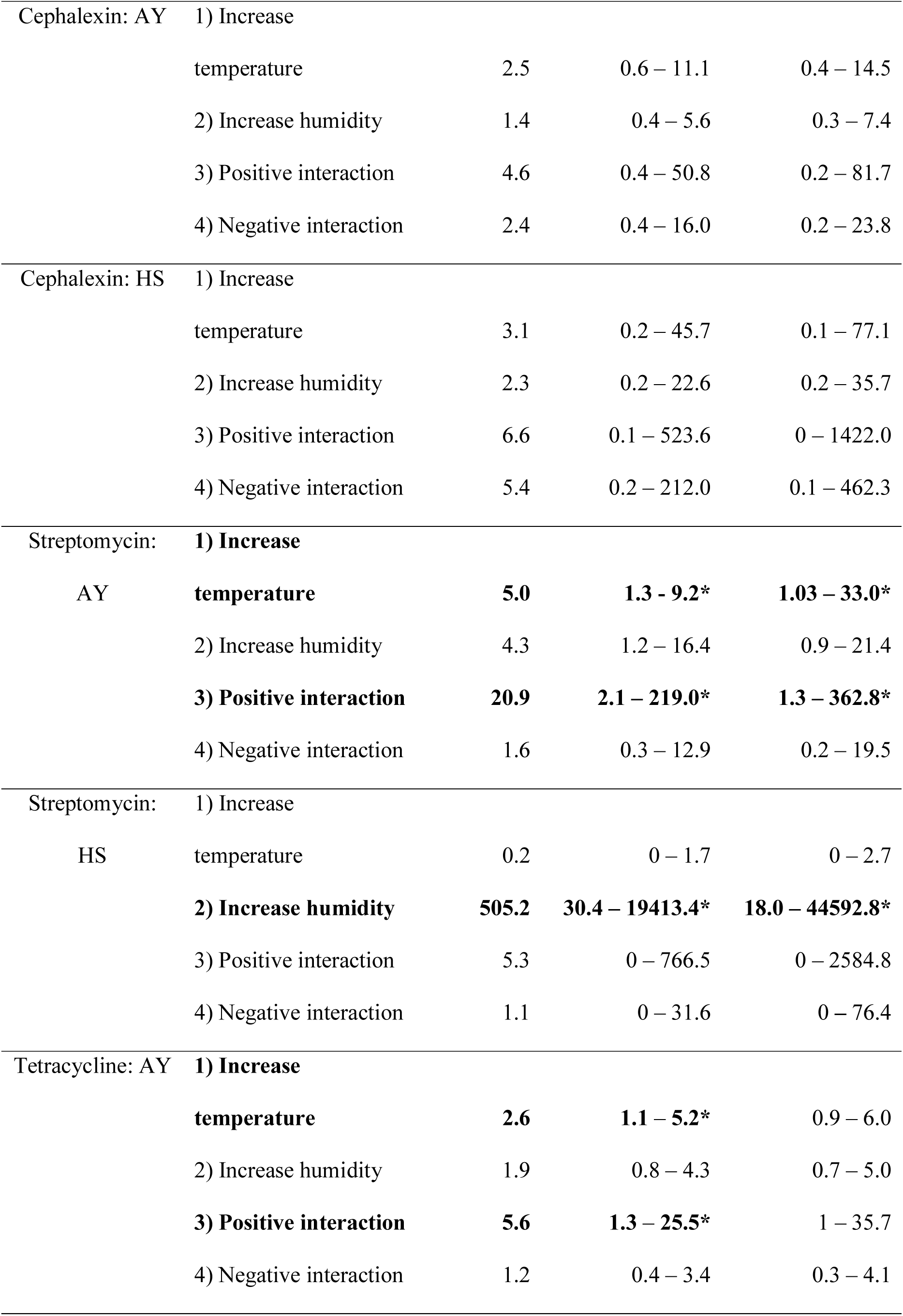
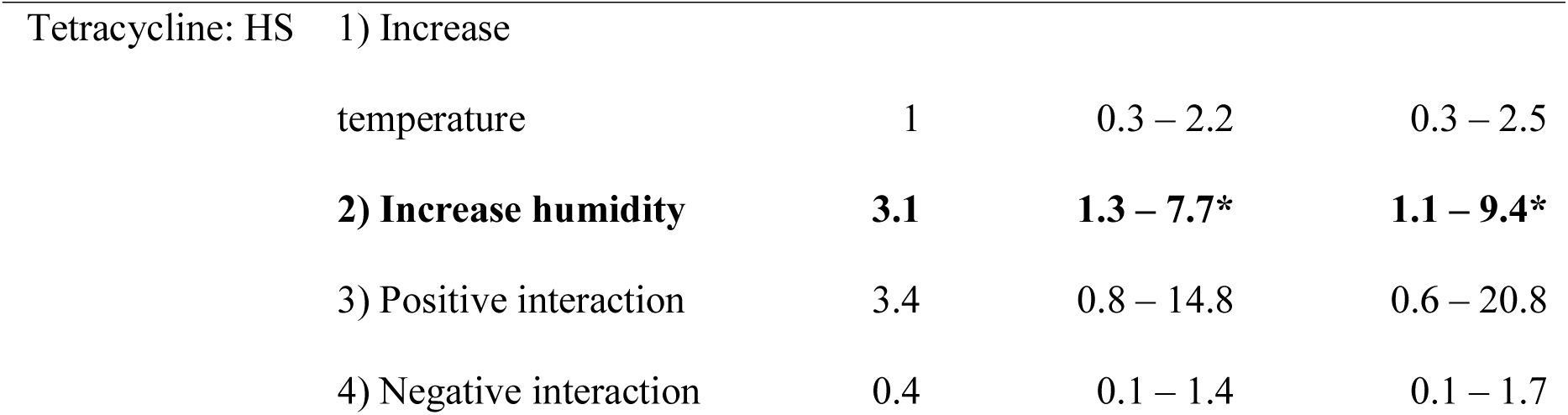
Change in model-estimated proportion of antimicrobial-resistant E. coli across weather condition change scenarios. Proportion change calculated from the model-estimated proportion of antimicrobial-resistant E. coli across four weather condition change scenarios: *1) ‘Increase temperature’: Increase from 1°C to 21°C, at 82% humidity* *2) ‘Increase humidity’: Increase from 60% to 92% humidity, at 10°C* *3) ‘Positive interaction’: From cold & dry (1°C, 60%) to hot & humid (21°C, 92%)* *4) ‘Negative interaction’: From cold & humid (1°C, 92%) to hot & dry (21°C, 60%)* *\* Credible interval does not cross the null (1)*

| Model | Weather condition<br>change scenarios | Proportion change in model-estimated<br>proportion of resistant isolates |  |  |
| --- | --- | --- | --- | --- |
|  |  | Median | 90% credible | 95% credible |
|  |  |  | interval | interval |
| Amoxicillin: AY | 1) Increase<br>temperature | 2.4 | 0.9 – 6.1 | 0.7 – 7.3 |
|  | **2) Increase humidity** | **3.4** | **1.3 – 10.5\*** | **1.1 – 14.7\*** |
|  | 3) Positive interaction | 5.6 | 0.9 – 29.1 | 0.6 – 41.5 |
|  | 4) Negative interaction | 0.8 | 0.2 – 2.8 | 0.1 – 3.6 |
| Amoxicillin: HS | 1) Increase<br>temperature | 2.5 | 0.7 – 7.1 | 0.6 – 8.5 |
|  | **2) Increase humidity** | **4.7** | **1.7 – 12.8\*** | **1.4 – 15.7\*** |
|  | **3) Positive interaction** | **16.0** | **2.6 – 104.9\*** | **1.7 – 162.4\*** |
|  | 4) Negative interaction | 0.7 | 0.1 – 3.0 | 0.1 – 3.9 |
| Cephalexin: AY | 1) Increase |  |  |  |
|  | temperature | 2.5 | 0.6 – 11.1 | 0.4 – 14.5 |
|  | 2) Increase humidity | 1.4 | 0.4 – 5.6 | 0.3 – 7.4 |
|  | 3) Positive interaction | 4.6 | 0.4 – 50.8 | 0.2 – 81.7 |
|  | 4) Negative interaction | 2.4 | 0.4 – 16.0 | 0.2 – 23.8 |
| Cephalexin: HS | 1) Increase |  |  |  |
|  | temperature | 3.1 | 0.2 – 45.7 | 0.1 – 77.1 |
|  | 2) Increase humidity | 2.3 | 0.2 – 22.6 | 0.2 – 35.7 |
|  | 3) Positive interaction | 6.6 | 0.1 – 523.6 | 0 – 1422.0 |
|  | 4) Negative interaction | 5.4 | 0.2 – 212.0 | 0.1 – 462.3 |
| Streptomycin: AY | **1) Increase** |  |  |  |
|  | **temperature** | **5.0** | **1.3 - 9.2\*** | **1.03 – 33.0\*** |
|  | 2) Increase humidity | 4.3 | 1.2 – 16.4 | 0.9 – 21.4 |
|  | **3) Positive interaction** | **20.9** | **2.1 – 219.0\*** | **1.3 – 362.8\*** |
|  | 4) Negative interaction | 1.6 | 0.3 – 12.9 | 0.2 – 19.5 |
| Streptomycin: HS | 1) Increase |  |  |  |
|  | temperature | 0.2 | 0 – 1.7 | 0 – 2.7 |
|  | **2) Increase humidity** | **505.2** | **30.4 – 19413.4\*** | **18.0 – 44592.8\*** |
|  | 3) Positive interaction | 5.3 | 0 – 766.5 | 0 – 2584.8 |
|  | 4) Negative interaction | 1.1 | 0 – 31.6 | 0 – 76.4 |
| Tetracycline: AY | **1) Increase** |  |  |  |
|  | **temperature** | **2.6** | **1.1 – 5.2\*** | 0.9 – 6.0 |
|  | 2) Increase humidity | 1.9 | 0.8 – 4.3 | 0.7 – 5.0 |
|  | **3) Positive interaction** | **5.6** | **1.3 – 25.5\*** | 1 – 35.7 |
|  | 4) Negative interaction | 1.2 | 0.4 – 3.4 | 0.3 – 4.1 |
| Tetracycline: HS | 1) Increase |  |  |  |
|  | temperature | 1 | 0.3 – 2.2 | 0.3 – 2.5 |
|  | **2) Increase humidity** | **3.1** | **1.3 – 7.7\*** | **1.1 – 9.4\*** |
|  | 3) Positive interaction | 3.4 | 0.8 – 14.8 | 0.6 – 20.8 |
|  | 4) Negative interaction | 0.4 | 0.1 – 1.4 | 0.1 – 1.7 |

**Figure S10:**
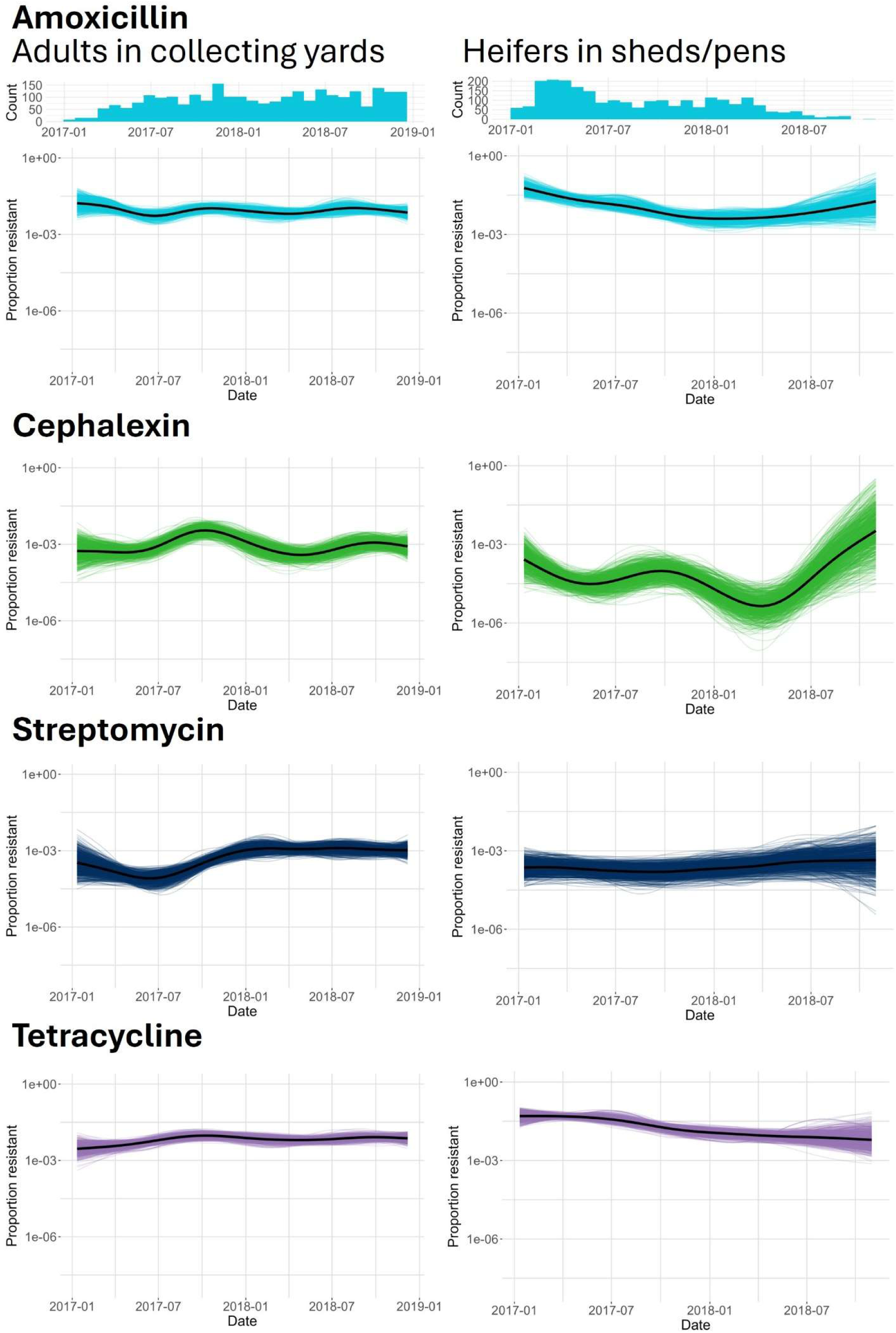
Estimated splines from non-linear models showing seasonal variation in the proportion of antimicrobial-resistant *E. coli* unaccounted for by the inclusion of weather-related covariates for each of the four tested antimicrobials across the two farm environments. Splines were estimated from 800 posterior draws and visualised as spaghetti plots. Thick black lines show median model-estimated proportions of antimicrobial-resistant E. coli across a range of observed temperature and humidity values, estimated from all 40000 posterior draws. For all splines the Y-axis shows log_10_ transformed proportion of antimicrobial-resistant E. coli. Histograms above splines show the distribution of samples across the sampling period, which are identical across antimicrobials, but vary between HS and AY groups.

